# Human follicular CD4 T cell function is defined by specific molecular, positional and TCR dynamic signatures

**DOI:** 10.1101/2020.05.12.089706

**Authors:** Kartika Padhan, Eirini Moysi, Alessandra Noto, Alexander Chassiakos, Khader Ghneim, Sanjana Shah, Vasilis Papaioannou, Giulia Fabozzi, David Ambrozak, Antigoni Poultsidi, Maria Ioannou, Craig Fenwik, Samuel Darko, Daniel C. Douek, Rafick-Pierre Sekaly, Giuseppe Pantaleo, Richard A. Koup, Constantinos Petrovas

## Abstract

The orchestrated interaction between follicular helper CD4 T cells (TFH) and germinal center (GC) B cells is crucial for optimal humoral immunity. However, the regulatory mechanisms behind spatial distribution and function of TFH is not well understood. Here, we studied human TFH cells and found that transitioning to a CD57^hi^ TFH status was associated with distinct positioning in the GC, phenotype, transcriptional signatures, function and downregulation of their T-cell receptor (TCR). Single cell TCR clonotype analysis indicated a unidirectional transition towards the CD57^hi^ TFH status, which was marked with drastic changes in the nature of immunological synapse formation where peripheral microclusters become dominant. Lack of central supra molecular activation cluster (cSMAC) formation in TFH synapse was associated with enhanced ubiquitination/proteasome activity in these cells. Our data reveal significant aspects of the tissue organization and heterogeneity of follicular adaptive immunity and suggest that CD57^hi^ TFH cells are endowed with distinctive programming and spatial positioning for optimal GC B cell help.

**One Sentence Summary:** human TFH cell heterogeneity

## Introduction

The development of immunogen and pathogen specific B cell responses requires the coordinated function of highly differentiated immune cell populations residing in the follicular and GC areas (1, 2). TFH cells provide critical help to B cells, particularly in GCs, that governs the fate of B cells by regulating the process of affinity maturation, class switching, somatic hyper-mutation and generation of memory B cell responses (3–5). This help is mainly mediated by surface receptors (1) and cytokines including IL-4 and IL-21 (6). TFH cells are characterized by a unique phenotype, gene signature and functional profile where many of their characteristics are preserved across different species (mouse, non-human primates, and human) (7–10).

TFH cell development is a multi-stage process (8, 11). Each stage is driven by a combination of factors including TCR stimulation, co-stimulation/co-inhibition, cytokines, chemokines and transcription factors (5, 12). The outcome of this differentiation process can presumably result in a highly heterogeneous pool with particular TFH cell subsets characterized by distinct profile and function (13–16). The fate of TFH cells after their peak during an immune response is not clear. It has been proposed that the expansion of TFH is followed by cell death or conversion to central memory CD4 T cells (17). Alternatively, TFH cells that survive could become memory TFH cells that can traffic to blood (18) or peripheral tissues; however, these cells do not express classic TFH cell biological markers (19).

Several studies have shown the role of receptors like PD-1 (20), CXCR5 (21) and ICOS (20) in TFH cell positioning in GC and function. Furthermore, modulation of TFH immunological synapse (IS) dynamics by neurotransmitters like dopamine (22) could affect TFH helper function. However, the connection between differentiation, spatial distribution, phenotype and the quality of TCR-induced signaling in TFH cells is not well understood. Our single cell analysis revealed a distinct phenotype, gene signature, spatial distribution and IS formation for two subsets (PD-1^hi^CD57^lo^ and PD-1^hi^CD57^hi^) of TFH cells. This heterogeneity characterized TFH cells in both tonsil and lymph nodes.

## Results

### Preferential localization of CD57^hi^PD-1^hi^ TFH cells in proximity to proliferating GC B cells, FDC network and GC dark zone

First we investigated the TFH heterogeneity by assessing the localization of TFH subsets, identified by the expression of PD-1 and CD57 (13–15). The intrafollicular localization of CD57^hi^PD-1^hi^ and CD57^lo^PD-1^hi^ TFH cells was investigated using a multiplexed confocal imaging assay (Figure 1A). Follicles were defined as CD20^hi/dim^, non-GC follicular areas as CD20^dim^Ki67^lo^, while dark zones were highly enriched in CD20^dim^Ki67^hi^ and light zones were populated mainly by CD20^hi^Ki67^lo/hi^ B cells, CD4 T cells and FDCs (Figure 1, A and B). CD57^hi^PD-1^hi^ TFH cells were closer to DZ compared to CD57^lo^PD-1^hi^ TF cells (Figure 1, A and B). 3D imaging of individual follicles further confirmed the preferential localization of CD57^hi^PD-1^hi^ TFH cells in proximity to DZ compared to CD57^lo^PD-1^hi^ TF cells (Figure 1C, Supplemental Figure 1A and movies 1, 2 and 3). A computational method for the quantitation of distances between follicular cell subsets or between cells and follicular areas (Supplemental Figure 1B) was performed at individual imaged tissue sections. To minimize inconsistencies due to tissue orientation, only follicles with well-defined, distinct intrafollicular areas were analyzed. A significantly shorter (longer) average distance between DZ (non-GC follicular area) and CD57^hi^PD-1^hi^ compared to CD57^lo^PD-1^hi^ TFH cells was found in each follicle tested (Figure 1D and Supplemental Figure 1C). A different distance to DZ between CD57^lo^PD-1^hi^Ki67^hi^ and CD57^lo^PD-1^hi^Ki67^lo^ was found too (Figure 1E). A similar profile was observed when the distance between TFH subsets and LZ total B cells (data not shown), Ki67^hi^CD20^hi/dim^ B cells or CD68^hi^ follicular macrophages was analyzed (Figure 1, F and G). A simultaneous analysis of distances between TFH subsets and DZ area, FDC network (Supplemental Figure 1D) and B cells, revealed a distinct positioning between the two TFH subsets with CD57^hi^PD-1^hi^ been closer to all other elements (Figure 1H). Therefore, CD57^hi^PD-1^hi^ TFH cells are characterized by a distinct spatial positioning that could promote the interaction between FDC, GC B and TFH cells.

**Figure 1.**
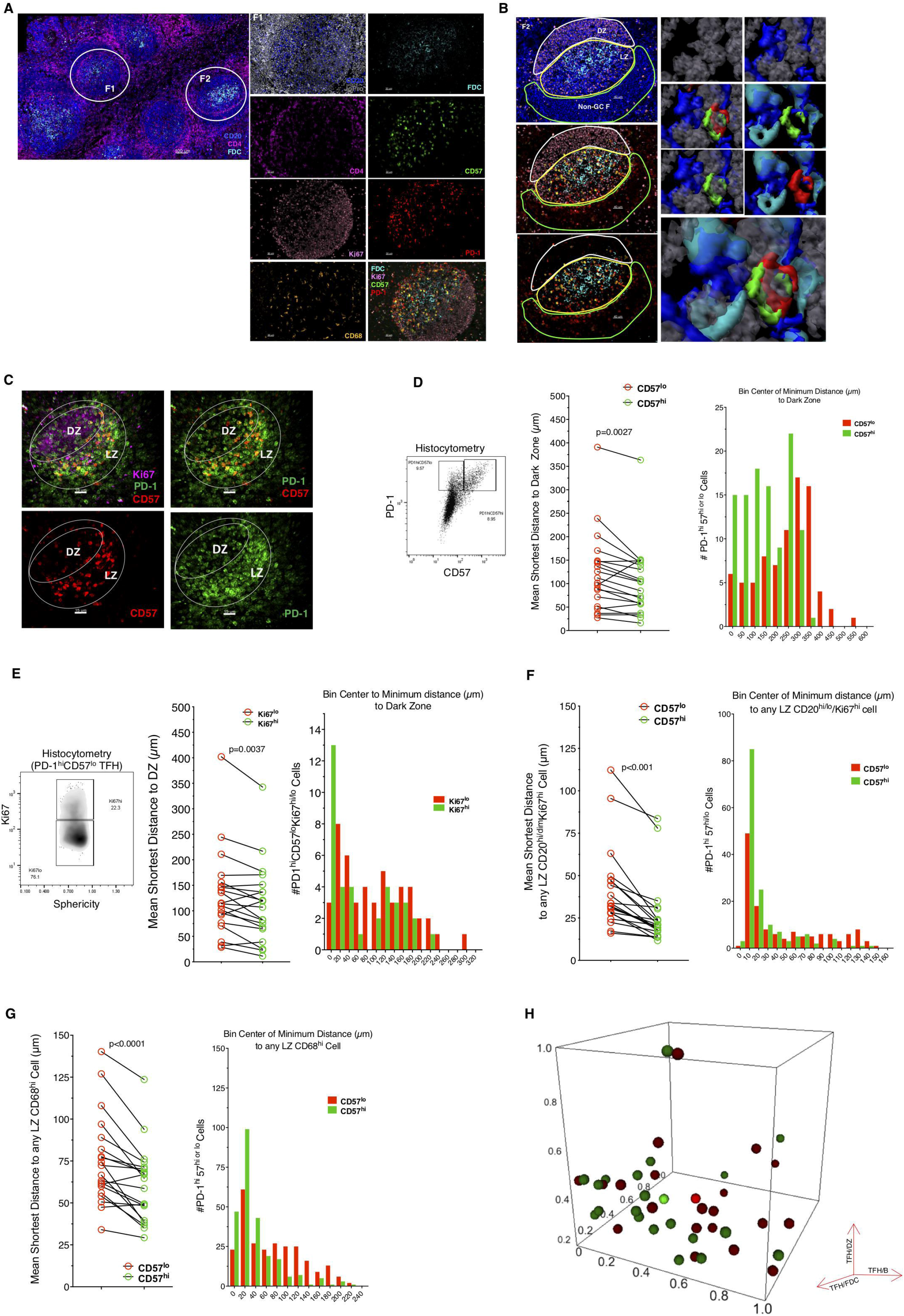
Preferential localization of CD57hi TFH cells proximal to Ki67hi GC B cells, FDC and GC-dark zone. (A) Representative confocal image (40X, scale bar: 100 mm) showing the expression of CD20, CD4 and FDC in a tonsillar section. Follicular area (F1) (scale bar: 30 mm) depicting single markers and two merged (CD20/JoJo and FDC/Ki67/CD57/PD-1) images are shown too. (B) A zoomed follicle (F2) (scale bar: 40 mm) and follicular areas as defined by the expression of relevant markers are shown; DZ (solid white line), LZ (solid yellow line) and Non-GC F (solid green line). Surface analysis showing different combinations of the volumetric expression of nuclear, CD20, CD57, PD-1 and FDC in F2. (C) The 2D projected localization of Ki67, CD57 and PD-1 cells of a 3D imaged follicle (supplementary video follicle 1) is shown. (D) TFH subsets (Histocytometry) and the mean shortest distance of CD57lo and CD57hi TFH cells to DZ are shown (n=20 follicles from 4 tonsils). Representative distribution histogram of CD57lo and CD57hi TFH cells with respect to their distance from the DZ in one follicle. (E) Ki67hi or lo CD57lo TFH cell subsets, their mean shortest distance to DZ and a representative distribution histogram are shown. The mean shortest distance of CD57lo and CD57hi TFH cells to CD20hi/dimKi67hi B cells (F) or CD68hi follicular cells (G) are shown (n=20 follicles from 4 tonsils) and relevant distribution histograms are shown. (H) The mean shortest distance (max/min normalized axis) between CD57hi or CD57lo TFH subsets and Ki67hi GC B cells, FDC and DZ is shown. Each sphere represents one follicle (CD57hiPD1hi -dark green, CD57loPD1hi-dark red). The mean value is also shown for CD57hiPD1hi (light green) or CD57loPD1hi (light red) TFH cell subsets. The Wilcoxon test was used for the analysis of data showed in D, E, F, and G.

### Unidirectional transition form CD57^lo^PD-1^hi^ to CD57^hi^PD-1^hi^ status is associated with distinct phenotypic, transcriptional and functional profile

Flow-cytometry analysis (Figure 2A and Supplemental Figure 2A), showed significantly higher relative frequency (%) of CD57^lo^ to CD57^hi^ TFH cells in human tonsils (Figure 2B). A significant correlation was found between CD57^lo^ TFH and all CD4 T cell subsets (naïve-CD27^hi^CD45RO^lo^PD-1^lo^, non-TFH-CD27^hi^CD45RO^hi^PD-1^lo^ and pre-TFH-CD27^hi^CD45RO^hi^PD-1^dim^) but CD57^hi^ TFH cells (Supplemental Figure 2B). Significantly increased frequency and expression per cell, in the “receptor^hi^” subsets, (judged by MFI-Mean Fluorescence Intensity) was found in CD57^hi^ compared to CD57^lo^ TFH cells for several surface receptors (Supplemental Figure 2, C and D), transcription factors (TFs) Bcl-6 and c-Maf and the proliferation marker Ki67 (Supplemental Figure 2E). Therefore, the unique localization of tonsillar CD57^hi^ TFH cells is linked to a distinct expression of receptors and TFs (Figure 2C) important for the development and function of TFHs.

**Figure 2:**
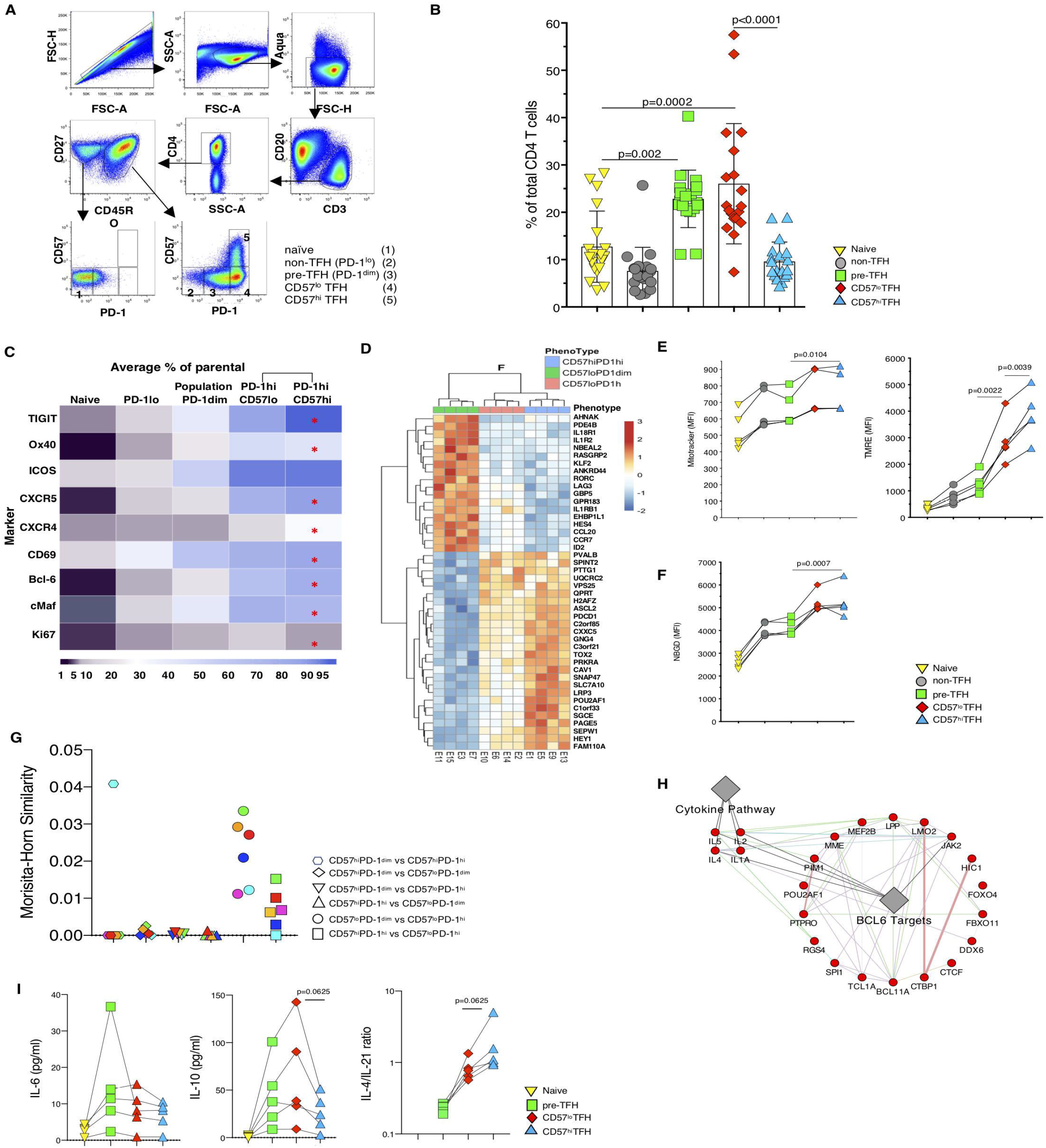
Tonsillar CD57hi and CD57lo TFH cells express distinct phenotypic, transcriptional and functional profiles. (A) Tonsillar CD4 T cell subsets identified by flow-cytometry. (B) Frequencies of CD4 T cell subsets, marked with different symbols (n=20 tonsils). Mean values (box height) and SD error bars are shown. (C) Relative expression of individual markers in tonsillar CD4 T cell subsets (* p<0.05). (D) Heatmap representing the expression of the top 50 variant genes based on an ANOVA (analysis of variance, F-test, p-val≤0.05) between CD57hiPD1hi (blue samples n=4), CD57loPD1hi (red samples, n=4) and CD57loPD1dim (green samples n=4). Rows represent genes and columns represent samples. Red and blue correspond to up- and down-regulated genes respectively. (E) *Ex vivo* levels of Mitotracker GreenFM and TMRE in tonsillar CD4 T cell subsets (n=5 tonsils). (F) Levels of *ex vivo* NBDG uptake by tonsillar CD4 T cell subsets (n=5 tonsils). (G) Single cell TCR clonotype analysis. The TCR repertoire overlap by cell subsets is shown (n=6 tonsils). (H) GSEA was used to assess the enrichment of cytokine signaling pathways (Reactome, KEGG c2, MSigDB) and BCL6 Targets (GeneRIF, TRANSFAC) between CD57loPD1hi and CD57hiPD1hi sorted populations. Circular nodes represent genes and edges reflect the association between these features. Color of the edge highlights the association between nodes. Red nodes indicate these genes are upregulated in CD57hiPD1hi compared to CD57loPD1hi. Biological annotation (diamond nodes) reflects function of genes. (I) The levels of secreted cytokines after *in vitro* stimulation of sorted CD4 T cell subsets are shown. The Wilcoxon test was used for the analysis of data showed in B, C, E, F and I. *p<0.006.

A distinct transcriptional profile was found between CD57^hi or lo^ TFH cells and CD57^hi or lo^ PD-1^dim^ CD4 T cells (Supplemental Figure 2F) with several genes differentially expressed between CD57^lo^PD-1^dim^, CD57^lo^PD-1^hi^ and CD57^hi^PD-1^hi^ cell subsets (Figure 2D, Supplemental Figure 2G and online supplemental information). Transition to CD57^lo^ TFH status was associated with significant upregulation of mitochondrial/metabolic pathways and downregulation of IFNγ related pathways (Supplemental Figure 3A) while a downregulation of metabolic pathways was found in CD57^hi^ compared to CD57^lo^ TFH cells (Supplemental Figure 3A). Furthermore, a significant enrichment of TFH development pathways was found in CD57^hi^ compared to CD57^lo^ TFH cells (Supplemental Figure 3B) underlined by the higher expression of TOX2 (Supplemental Figure 2C), a positive regulator for TFH cell development (23). *Ex vivo* flow cytometry analysis (Supplemental Figure 3D), showed a significantly increased binding of TMRE (a marker for mitochondrial membrane potential (24)) and Mitotracker green FM (a surrogate for “mitochondrial mass” (25)) in TFH compared to non/pre-TFH cells (Figure 2E), consistent with their pathway signatures (Supplemental Figure 3A). A significant difference was also found for TMRE binding between CD57^lo^ and CD57^hi^ TFH cells (Figure 2E). Use of NBDG (Supplemental Figure 3D), a nondegradable glucose analogue (26), showed increased glucose uptake by TFH cell subsets (Figure 2F), in line with their molecular pathway profile (Supplemental Figure 3A). In line with our transcriptional data, significantly higher levels of c-myc and inducible expression of p53 after treatment with 50 μM etoposide (27), was found in TFH vs non/pre-TFH cells, particularly in CD57^lo^ TFH cells (Supplemental Figure 3, E-F). Single cell TCR clonotype analysis was carried out using the 10x Chromium sequencing platform. Our bulk gene array data, from sorted tonsillar CD4 subsets, were used to assign each single cell derived transcriptome to a particular CD4 subset (Supplemental Figure 3G). A significant TCR repertoire overlap was found only within the CD57^lo^PD-1^dim^ / CD57^lo^PD-1^hi^ and CD57^hi^PD-1^hi^ / CD57^lo^PD-1^hi^ comparisons (Figure 2G).

Further transcriptional analysis showed increased expression of i) cytokines like IL-4, IL-5 and IL-1A, and ii) several downstream targets of Bcl-6, in line with its expression (Supplemental Figure 2E), in CD57^hi^ compared to CD57^lo^ TFH cells (Figure 2H). *In vitro* stimulation of sorted tonsillar CD4 subsets showed a significantly lower secretion of Th1 type cytokines by TFH cells (Supplemental Figure 3H). Recently, a progressive differentiation of TFH cells, characterized by lower production of IL-21 and increased secretion of IL-4, was found to be associated with promotion of antibody secreting B cells (11). We found lower IL-10 secretion and a shift from IL-21 to IL-4 production by CD57^hi^ compared to CD57^lo^ TF cells (Figure 2I and Supplemental Figure 3H). Hence, the CD57^hi^ TFH cell unique GC localization is associated with distinct molecular, metabolic and functional profile.

### Significant reduction of TCR in TFH cells, particularly the CD57^hi^ ones

Imaging (Figure 3A) and histocytometry analysis (Figure 3, B and C) consistently revealed dimmer expression of CD3, but not CD27, in the follicular compared to extrafollicular areas. Flow cytometry (Supplemental Figure 4A) verified this profile in two separated experiments. A significant reduction of surface CD3 levels (MFI) was found in TFH cells, especially the CD57^hi^ ones, for both TCR chains tested (Figure 3D and Supplemental Figure 4B). CD150(SLAM)^lo^ TFH cells exhibit a distinct functional and *in vivo* cycling profile compared to CD150^hi^ TFH cells (8). The % of CD150^hi^ cells was significantly reduced in TFH cells, especially the CD57^hi^ ones (Figure 3E and Supplemental Figure 4, C and D). Absence of CD150 was associated with further reduction of TCR in the populations tested (Figure 3F and Supplemental Figure 4D). In line with this profile, several TCR-signaling genes were found differentially expressed between PD-1^dim^ and PD-1^hi^CD57^lo^ TFH cells (Figure 3G) and between PD-1^hi^CD57^lo^ and PD-1^hi^CD57^hi^ TFH cells (Supplemental Figure 4E). Thus, TFH differentiation is associated with major alterations related to TCR levels and possibly TCR-induced signaling.

**Figure 3:**
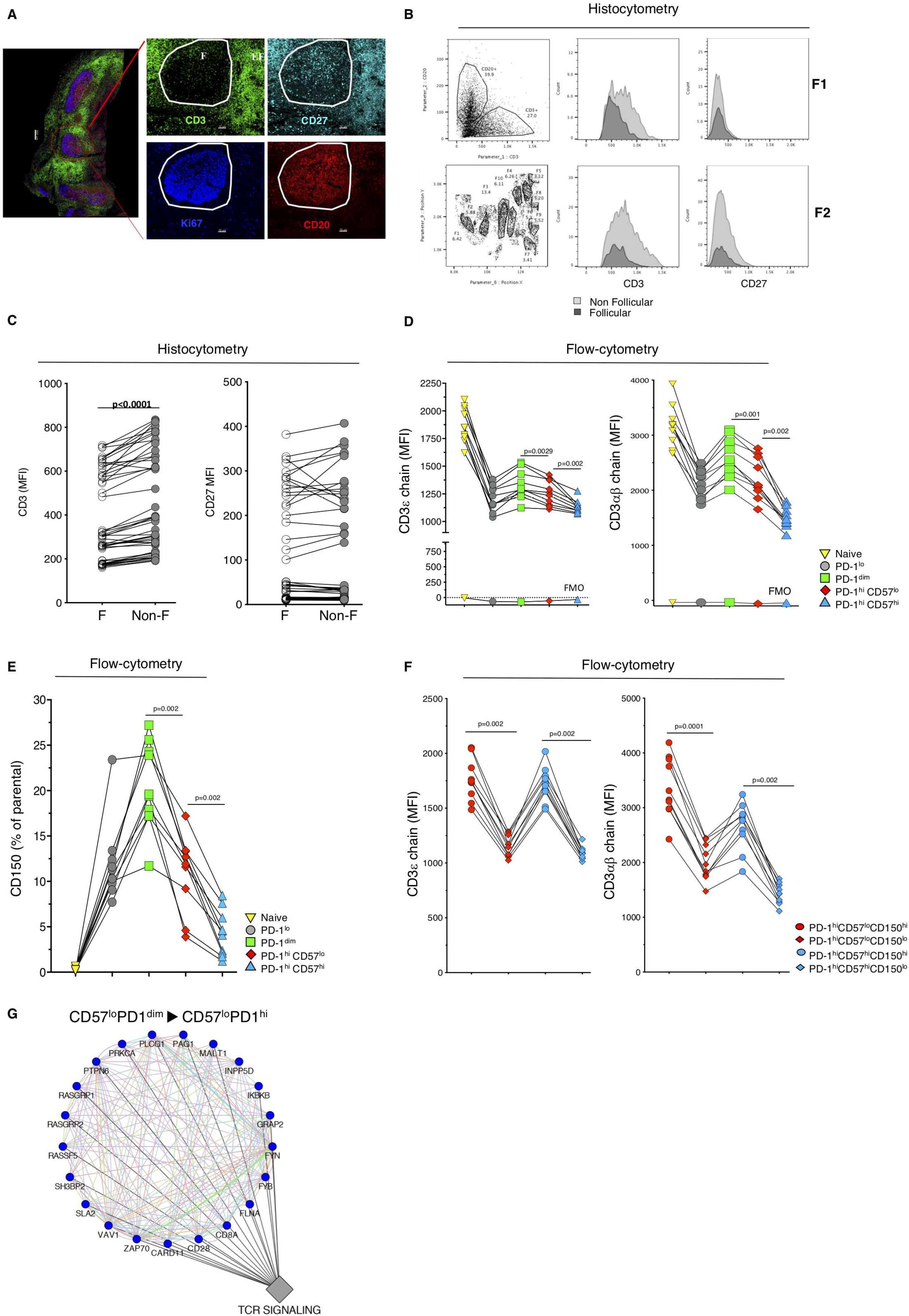
Differentiation towards the CD57hi TFH stage is associated with significant reduction of TCR. **(A)** Confocal image (40x, scale bar: 200 mm) showing the localization of CD3 (green), CD20 (red) and Ki67 (blue) positive cells in a tonsil tissue section (upper panel). A zoomed follicular area (red circle) (scale bar: 70 mm) with individual markers is shown too (white circle marks the borders of the follicle). **(B)** Histocytometry-generated plots showing the CD20 B and CD3 T cell populations, the identified follicular areas as well as histograms depicting the expression (intensity, MFI) of CD3 and CD27 in two non-follicular (gray) and follicular (dark gray) areas. **(C)** Graphs showing accumulated data of CD3 and CD27 MFI in follicular (F) and non-follicular (Non-F) areas (n=48 follicles). **(D)** Graphs showing the expression of CD3e (left) and CD3ab (right) in tonsillar CD4 T cell subsets marked with different symbols (n=10 tonsils). The corresponding FMO levels are also shown for comparison. **(E)** Graph showing the relative frequency of CD150hi cells in different CD4 T cell subsets. **(F)** graphs showing the expression of CD3e (left) and CD3ab (right) in tonsillar CD150hi or lo CD4 TFH cell subsets (n=10 tonsils). **(G)** GSEA was used to assess the enrichment of TCR signaling pathways (PID c2, MSigDB) between CD57loPD1hi and CD57loPD1dim sorted populations. A co-expression network of the leading-edge genes of TCR signaling was plotted to highlight their interaction. Gene Mania algorithm was used to infer network connections and co-expression. Circular nodes represent genes and edges reflect the association between these features. Color of the edge highlights the association between nodes. Blue nodes indicate these genes are down regulated in CD57loPD1hi compared to CD57loPD1dim. Biological annotation (diamond nodes) reflects function of genes.

### Human lymph node and tonsillar TFH cells share similar localization, phenotype and TCR downregulation profile

LN-derived cells from healthy, non-hyperplastic LNs were analyzed using a Cytof assay (Supplemental Figure 5A). In contrast to tonsils, TFH cells represent a small fraction of LN cells (Figure 4A). LN sections were analyzed by a multiplexed imaging assay where several active follicles were identified based on the presence of relevant populations (FDC, TFH) (Figure 4B and Supplemental Figure 5B). Quantitative imaging analysis showed that similar to tonsils, CD57^hi^PD-1^hi^ LN TFH cells were found closer to the DZ and proliferating GC B cells (Figure 4C and Supplemental Figure 5C). Several of the investigated factors were significantly upregulated in LN TFH cells (Supplemental Figure 5, D and E). Furthermore, CD57^lo^ (cluster 1) and CD57^hi^ (cluster 7) LN TFH cells exhibit distinct phenotypes (Figure 4, D and E and Supplemental Figure 5, F and G). Importantly, TFH differentiation was associated with significant reduction of CD3ε and CD3αβ (Figure 4F), especially in the absence of CD150 (Figure 4G and Supplemental Figure 5H). Therefore, tonsillar and LN TFH cells share a similar tissue localization, phenotype and TCR dynamics, justifying the use of tonsillar TFH for further analysis of TCR downregulation in TFH biology.

**Figure 4:**
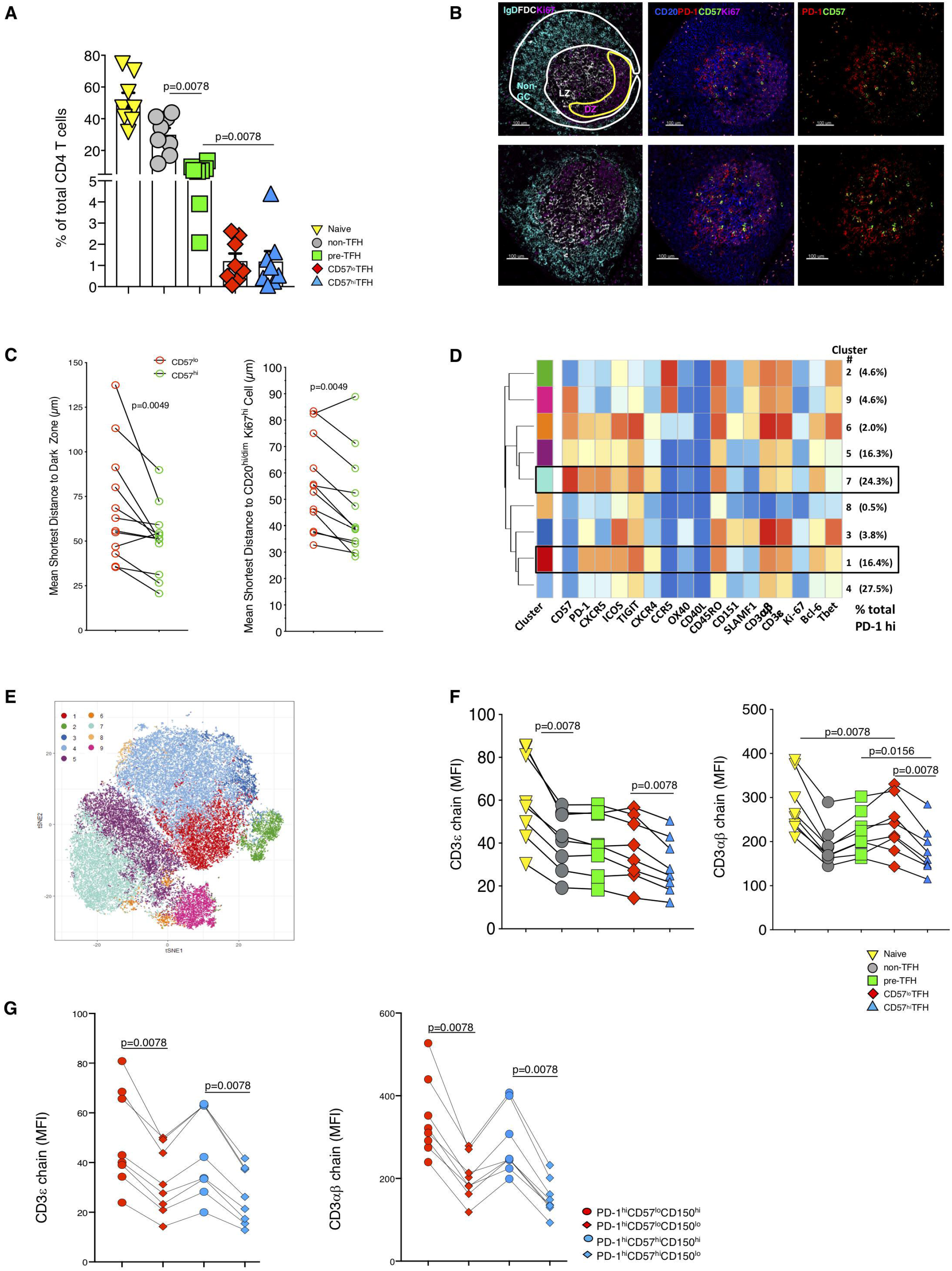
Human lymph node and tonsillar TFH cells express similar profiles. **(A)** Relative frequencies of individual CD4 T cell subsets, marked with different symbols, in human LNs (n=8). Mean values (box height) and SD error bars are shown. **(B)** Confocal images (scale bar: 100 um) showing the localization of IgDhi (cyan), FDChi (white), Ki67hi (magenta), CD20hi/dim (blue), PD-1hi (red) and CD57hi (green) cells in two follicular areas from a healthy LN. Solid lines depict the borders for particular follicular areas (white: Non-GC, yellow: GC-DZ). **(C)** Graphs showing the mean shortest distance of CD57lo (green) and CD57hi (red) TFH cells to DZ (left panel) and CD20hi/dimKi67hi B cells in 12 follicles (n=5 LNs). **(D)** Heat map showing 9 clusters identified based on the expression of 17 markers analyzed (showed at the bottom of the heat map) in total PD-1hi CD4 T cells. Clusters 1 (16.4%) and 7 (24.3%), highlighted with a black box, correspond to the two major TFH cell populations (cluster 1/CD57lo and cluster 7/CD57hi). **(E)** tSNE analysis performed on the total PD-1hi CD4 T cells using the data from all eight LNs. **(F)** the expression of CD3e (left) and CD3ab (right) in LN CD4 T cell subsets marked with different symbols (upper panel) and in TFH subsets with respect to CD150 expression (**G)** (n=10 LNs analyzed). The Wilcoxon test was used for the analysis of data showed in C, F and G.

### Tonsillar TFH cells form immunological synapse lacking cSMAC

Next, we asked whether the observed TCR expression impacts IS formation. We found a higher expression of CD18 (integrin β-chain-2) in CD57^lo^ TFH compared to PD-1^dim^ and CD57^hi^ TFH cells while active LFA-1 was significantly higher in TFH compared to non/pre-TFH cells, particularly in CD57^hi^ TFH cells (Supplemental Figure 6, A and B). In line with the transcriptome data, CD151, a component of IS (28), was also found increased in CD57^hi^ TFH compared to non/pre-TFH cells and CD57^lo^ TFH cells (Supplemental Figure 5H and 6B). An *in silico* IS formation assay (29, 30) (Supplemental Figure 6C), using anti-CD3 (OKT3) for TCR stimulation at a concentration inducing sub-optimal activation judged by the expression of CD69 (Supplemental Figure 6D), was performed. We sorted TFH cells based on ICOS instead of PD-1 to exclude any interference from PD-1 mediated inhibitory effect. Likewise, we avoided use of anti-CD3 antibodies in the sorting to avoid TCR stimulation. For comparison, we also sorted TFH cells based on PD1 and CXCR5 expression in a control experiment and found that PD-1 and ICOS represent same populations of TFH in terms of phenotype and synapse formation. Sorted TFH (CXCR5^hi^ICOS^hi^), pre-TFH (CXCR5^dim^ICOS^dim^) and CXCR5^lo^ICOS^lo^ (enriched in naïve cells and a lesser amount of memory CXCR5^lo^ICOS^lo^ cells) were used (Supplemental Figure 6E and methods). CXCR5^lo^ICOS^lo^ cells formed an IS characterized by TCR sequestration in the center (central supra molecular activation cluster, cSMAC), surrounded by integrins (peripheral SMAC) (Figure 5A), a profile less frequent in pre-TFH and TFH cells (Figure 5A). By counting the number of cells (Supplemental Figure 7A), we found a significantly higher number of TFH cells forming pSMAC IS compared to pre-TFH and naïve T cells (Figure 5B). In line with this pattern, TFH cells, which express a higher content of actin, an important mediator of IS formation (31), formed IS associated with significantly larger “actin depleted zone” (Supplemental Figure 7B). TFH and pre-TFH cells had significantly increased recruitment of TCR microclusters and ICAM to the synapse compared to naïve cells (Supplemental Figure 7, C and D). Recording the process of IS formation in live cells, further confirmed that naïve cells sort peripheral TCR microclusters into the cSMAC while TFH cells generate peripheral TCR microclusters that fail to move to the center of synapse (movies 4, 5 and 6).

**Figure 5:**
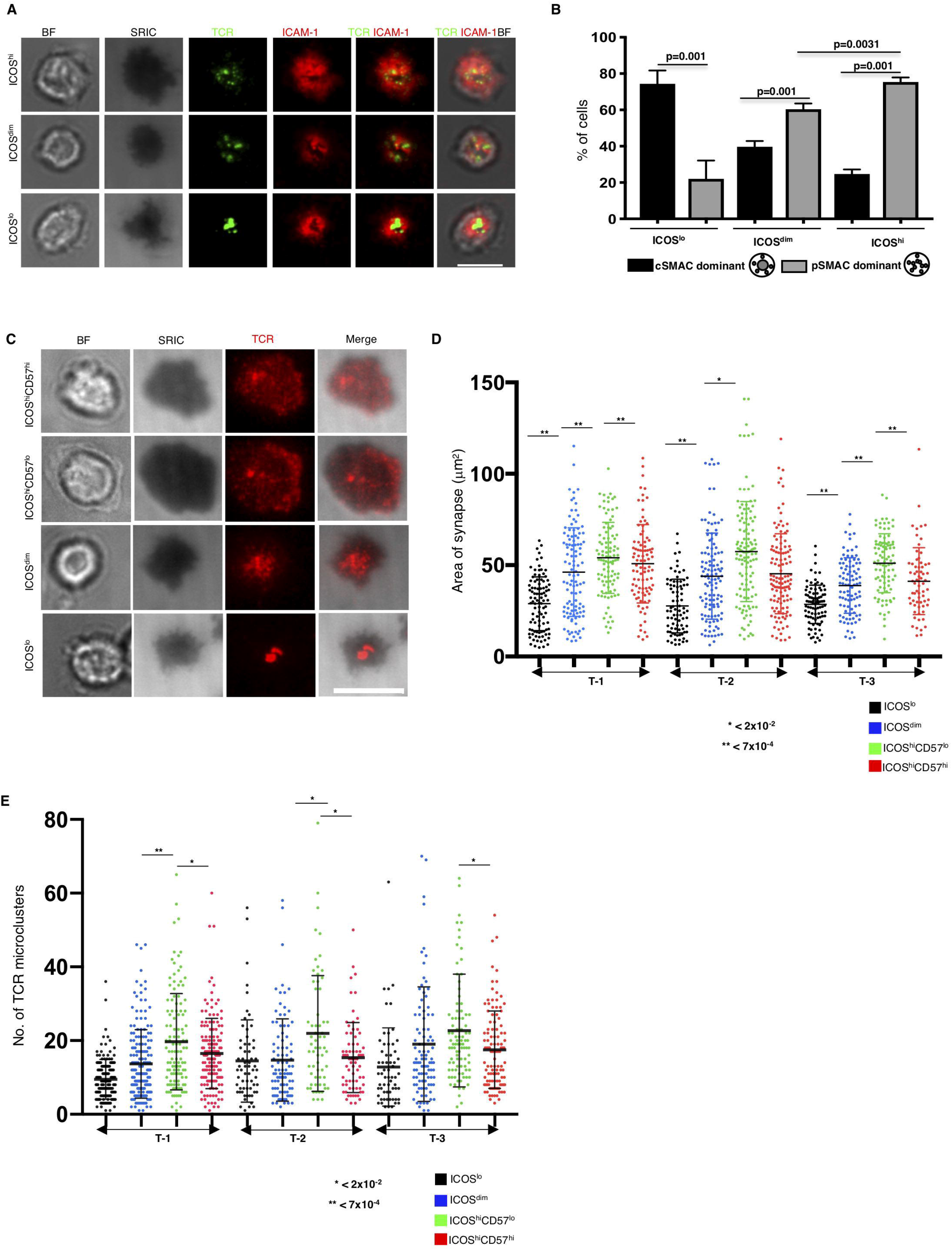
TFH cells form immunological synapse that lacks cSMAC. **(A)** Representative TIRF microscopy generated images (scale bar: 7 mm) showing the bright field (BF), surface reflection interference contrast (SRIC), TCR (green), ICAM-1 (red) as well as merged images of individual sorted tonsillar CD4 T cells expressing an ICOSlo, ICOSdim or ICOShi phenotype. **(B)** Bar graph depicting the frequency of cells forming a cSMAC or pSMAC type of IS in the ICOSlo, ICOSdim or ICOShi CD4 T cells (n=3 tonsils) (unpaired t test was used for statistical analysis). Mean values (horizontal lines) and SD bars are shown. **(C)** Representative TIRF microscopy generated images (scale bar: 10 mm) showing the BF, SRIC, TCR (red) as well as a merged image of individual sorted tonsillar CD4 T cells expressing an ICOSlo, ICOSdim, CD57loICOShi or CD57hiICOShi phenotype. **(D)** Dot plots showing the synaptic area measured by calculating the area of SRIC image (Welch ANOVA was used for statistical analysis). **(E)** The numbers of TCR microclusters in the synaptic area are shown (Kruskal-Wallis and Conover-Inman methods were used for statistical analysis). Sorted cells from three tonsils were used. Each dot represents an individual cell. Sorted populations are marked with different color (ICOSlo-black, ICOSdim-blue, CD57loICOShi-green and CD57hiICOShi-red). Mean values (horizontal lines) and SD bars are shown.

Next, the IS formation was investigated in sorted TFH subsets (Supplemental Figure 6E). Both TFH subsets formed a pSMAC IS (Figure 5C). The size of synapse was measured by quantifying the spread area in interference contrast image (Supplemental Figure 7A). CD57^lo^ TFH cells formed significantly largest synapse and recruited the greatest number of TCR microclusters to the synapse compared to naïve, pre-TFH and CD57^hi^ TFH cells (Figure 5, D and E) while lower average synaptic TCR intensity was found in CD57^hi^ compared to CD57^lo^ TFH cells (Supplemental Figure 7E). Hence, CD3 downregulation in TFH cells is associated with the formation of pSMAC-oriented IS.

### Peripheral clusters in CD57^hi^ TFH cell IS favor sustenance of TCR signaling

As TCR and associated molecules in the proximal signaling complex are tyrosine phosphorylated upon T cell activation, pTyr could be used as a read-out of early TCR-induced signaling. TFH cells accumulated TCR and pTyr at the peripheral IS clusters (Figure 6A). CD57^lo^ TFH cells had the highest number of IS pTyr microclusters (Figure 6B), while CD57^hi^ TFH cells had the lower average IS pTyr intensity (Figure 6C). Tissue imaging showed that total cellular pTyr was mainly found in the GC/LZ area with a “dotted” and a less frequent “polarized” pattern identified (Supplemental Figure 8A). The specificity of our assay was verified by staining tissues in the presence of λ-phosphatase (Supplemental Figure 8B). We found higher *in vivo* total cellular pTyr expression in CD57^hi^ compared to CD57^lo^ TFH cells (Figure 6D and Supplemental Figure 8C), confirming the competence for tyrosine phosphorylation in CD57^hi^ TFH cells.

**Figure 6:**
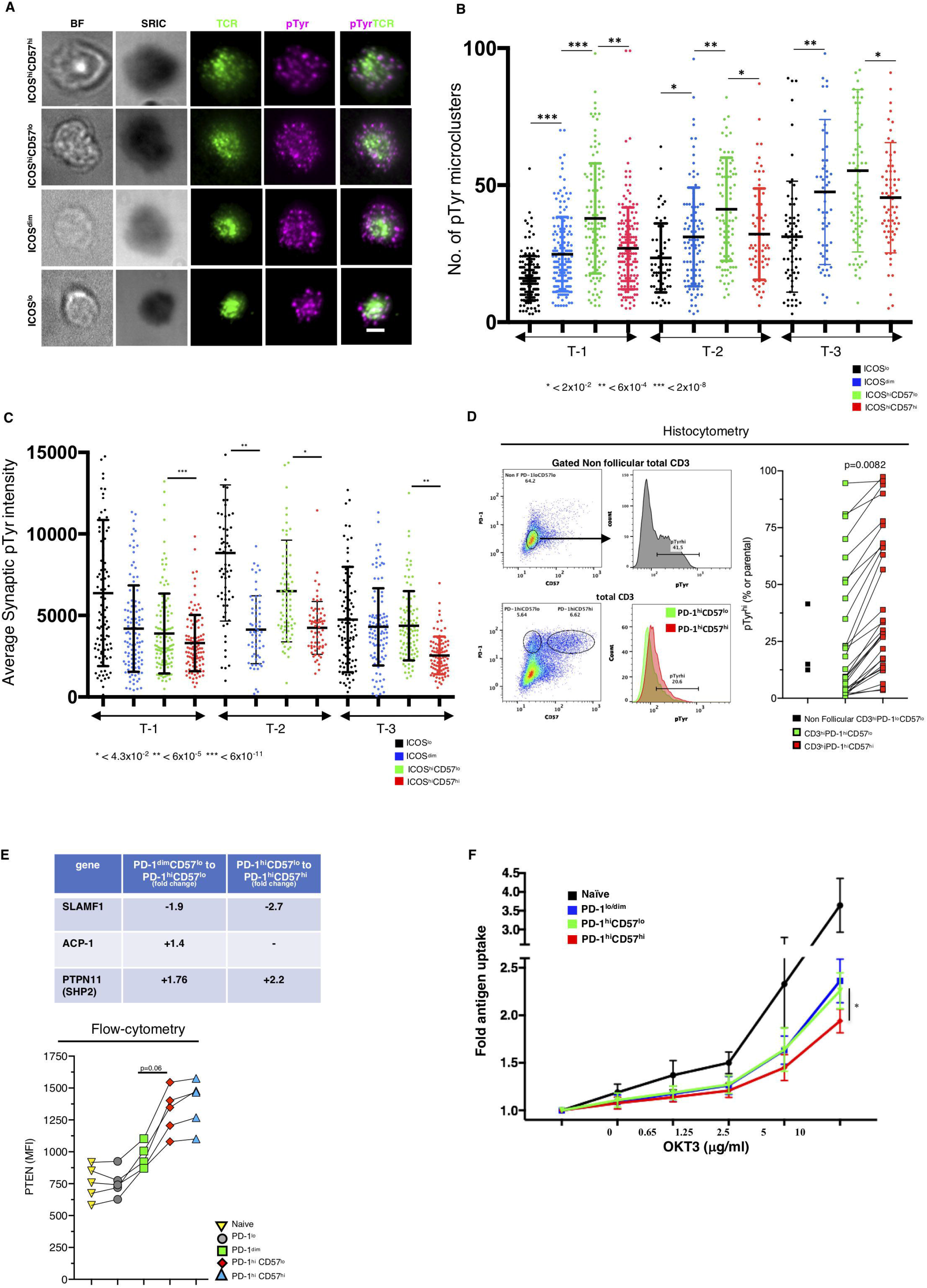
Altered early TCR-induced signaling in CD57hi TFH immunological synapse. **(A)** Representative TIRF microscopy generated images (scale bar: 6 um) showing the BF, SRIC, TCR (green), pTyr (magenta) as well as a merged image of individual sorted tonsillar CD4 T cells expressing an ICOSlo, ICOSdim, CD57loICOShi or CD57hiICOShi phenotype. (**B**) Accumulated data showing the synaptic number of pTyr microclusters. Mean values (horizontal lines) and SD bars are shown. (**C**) Accumulated data showing the average synaptic intensity of pTyr Sorted populations, marked with different color (ICOSlo-black, ICOSdim-blue, CD57loICOShi-green and CD57hiICOShi-red), from three tonsils were used. Each dot represents an individual cell. Welch ANOVA method was used for the statistical analysis. Mean values (horizontal lines) and SD bars are shown. **(D)** Histocytometry generated 2D plots showing the expression of PD-1 and CD57 in non-follicular and total CD3 T cells as well as the expression (MFI) or pTyr in marked subsets (left panel). Accumulated data depicting the relative frequency of pTyrhi in non-follicular CD3hiPD-1loCD57lo (black squares) and follicular CD3hiPD-1hiCD57lo (green squares) and CD3hiPD-1hiCD57hi (red squares) T cells. Each green and red square represents an individual follicle (n=29 follicles) from three tonsils were analyzed. **(E)** The relative expression of three phosphatase genes in indicated sorted tonsillar CD4 T cell subsets is shown (upper panel). The expression of PTEN (MFI) in different tonsillar CD4 T cell subsets (marked with different symbols) is shown (n=5 tonsils analyzed) (lower panel). Data were analyzed with the Wilcoxon test. **(F)** graph showing the dose dependent uptake (fold change over ICAM-1 control) of labeled anti-CD3 antibody (OKT3) by PDlo (black), PD-1dim (blue), CD57loPD-1hi (green) and CD57hiPD-1hi (red) tonsillar CD4 T cells (n=5 tonsils analyzed). The data were analyzed using student t test. SD bars are shown.

Increased expression of genes encoding for phosphatases related to TCR-induced signaling (ACP-1, SHP2) (32) and SLAMF1 (CD150), that interacts with SHP-2 (33) and could potentially interfere with the tyrosine phosphorylation of proteins, was found in TFH cells, especially the CD57^hi^ ones (Figure 6E). Furthermore, PTEN (phosphatase and tensin homolog), a negative regulator of T cell activation (34), was also higher in TFH cells (Figure 6E and Supplemental Figure 8D). Performance of an antigen uptake assay (Supplemental Figure 8E) showed that CD57^hi^ TFH cells had the lowest capability to uptake antigen (αCD3-Alexa488) (Figure 6F) indicating that the formation of pSMAC IS helps CD57^hi^ TFH cells display TCR and associated molecules in microclusters where compromised endocytosis of these molecules may be crucial for sustenance of signal *in vivo*.

### Lack of cSMAC formation in TFH cells is associated with increased proteasome activity

We sought to investigate possible mechanisms for the observed lack of cSMAC in TFH cells. The transition from pre-TFH to TFH was associated with increased expression of ubiquitination and proteasome related genes, critical biological factors for IS formation (35), (Figure 7A). TFH cells had high poly-ubiquitination level linked to reduced free ubiquitin (Figure 7B), significantly higher levels of Lysine-48 (K-48), that targets proteins for proteasomal degradation(36) (Figure 7C) and higher levels of *ex vivo* proteasomal activity compared to naïve and pre-TFH cells (Figure 7D and Supplemental Figure 8F). Blocking the proteasome activity with MG132 resulted in compromised formation of cSMAC and showed increased recruitment of TCR microclusters in the naive (Figure 7E) but not the TFH synapse (Supplemental Figure 8G). We hypothesize that the observed deficiency of free ubiquitin pool as a result of high proteasomal activity could contribute to the lack of cSMAC formation in TFH cells.

**Figure 7:**
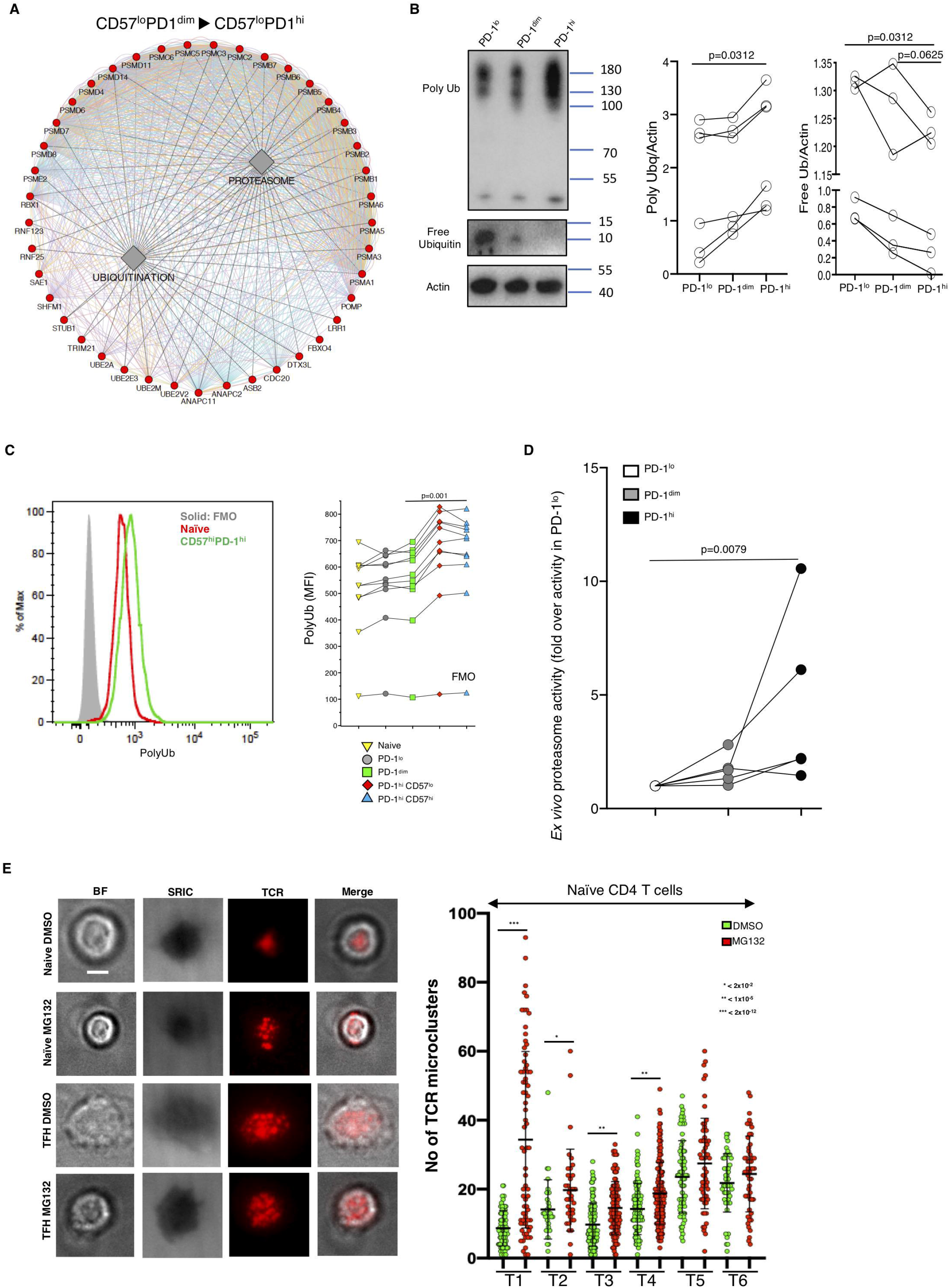
Lack of cSMAC in TFH immunological synapse is associated with increased ubiquitination and proteasome activity. **(A)** A co-expression network of the leading-edge genes of Proteasome/Ubiquitination pathways between CD57loPD1hi and CD57loPD1dim sorted populations. Circular nodes represent genes and edges reflect the association between these features. Color of the edge highlights the association between nodes. Red nodes indicate these genes are upregulated in CD57loPD1hi compared to CD57loPD1dim. Biological annotation (diamond nodes) reflects function of genes. **(B)** Western blot analysis of polyubiquitinylated proteins, free ubiquitin and actin in cell extracts from sorted nai□ve, pre- and TFH sorted tonsillar cells. Ratios of poly- or free-Ubq to actin in relevant cell extracts are shown (n=6 tonsils, Wilcoxon test for analysis). **(C)** Flow cytometry histogram showing the expression of polyubiquitinylated proteins in nai□ve and CD57hiPD-1hi TFH tonsillar cells (the FMO for the polyubiquitination antibody is shown too). The expression of polyubiquitinylated proteins in tonsillar CD4 T cell subsets is shown too (n=10 tonsils). Wilcoxon test was used for data analysis. **(D)** Graph showing the *ex vivo* proteasome activity (fold induction over the activity of PD-1lo cells) in sorted CD4 T cells expressing a PD-1lo, PD-1dim or PD-1hi phenotype (data were analyzed with Mann-Whitney test). **(E)** Representative TIRF microscopy generated images (scale bar: 6um) showing the bright field (BF), interference, TCR (red) and a merged image of individual sorted nai□ve or TFH cells (left panel). Sorted cells were treatment with DMSO (carrier) or MG132 10mM for 2hr before the incubation on the lipid bilayers. Dot plot showing the number of synaptic TCR microclusters in nai□ve tonsillar CD4 T cells (n=6 tonsils) treated with DMSO (green) or MG132 (red) (right panel). The Kruskal – Wallis and Conover - Inman methods were used for data analysis.

## Discussion

Spatial organization of relevant immune cells is of great importance for the optimum outcome of local tissue immunoreactions (37). The follicular architecture and the abundancy of relevant cell populations make tonsil an ideal organ to analyze TFH heterogeneity, particularly with respect to their spatial positioning. A compartmentalization of TFH cell subsets in the follicle/germinal center was found, characterized by the preferential proximity of CD57^hi^ TFH cells to FDC network, DZ and Ki67^hi^GC B cells. Our 3D imaging confirmed that this is a generalized profile applying across the follicle/GC (38). CD57^hi^ and Ki67^hi^CD57^lo^ TFH cell subsets were found in close proximity with DZ suggesting an *in situ*, within the GC, differentiation of dividing CD57^lo^ to CD57^hi^ TFH cells. The described spatial positioning suggests a higher possibility for CD57^hi^ TFH cells to support the multi-interaction between antigen, FDC, TFH and GC B cells for the delivery of optimal help to B cells.

This differential localization was also linked to distinct phenotypes, molecular signatures and function between the two TFH subsets studied. An altered *in vivo* turnover, supported by the different expression of Ki67, or further differentiation i.e. to memory TFH cells that possibly do not express typical TFH markers (19) could contribute to the lack of association found between CD57^lo^ and CD57^hi^ TFH cells. The differential expression of several receptors, and presumably the receptor-mediated signals, further contributes to the heterogeneity of the TFH pool. The transition from pre-TFH to CD57^lo^ TFH stage was characterized by a molecular profile of increased metabolism and mitochondria function. In line with previous studies showing that glycolysis promotes differentiation of TFH cells (12, 39) we found a significantly higher capacity for glucose uptake and possibly glycolytic function (upregulation of glycolysis pathway) in CD57^lo^ TFH compared to non/pre-TFH cells. Higher induction of p53, a pluripotent factor acting both at DNA repair and mitochondrial homeostasis level (40), could affect CD57^lo^ TFH cell development particularly in chronic diseases like HIV/SIV infection where the operation of such pathways could affect the TFH cell survival (6, 41) as well as the dynamics of associated virus (42). Compared to CD57^lo^, CD57^hi^ TFH cells were characterized by a potentially lower metabolic activity (downregulation of glycolysis and oxidative phosphorylation pathways) and a significantly higher mitochondrial membrane potential, a hyperpolarization stage that could represent a pre-apoptotic status (43). The lower capacity for upregulation of p53 in CD57^hi^ TFH cell could further compromised their response to stress conditions present in the follicular/germinal center areas. Thus, CD57^hi^ TH cells are characterized by a less metabolically active status and a lower capacity for division compared to CD57^lo^ TFH cells. Their closer proximity to CD68^hi^ follicular macrophages which uptake/phagocytose dying cells in situ (44), suggests a higher cell death rate compared to CD57^lo^ TFH cells. TCR repertoire analysis has been used for tracking distinct T cell subpopulations and possible “communication” between cellular compartments (45). Measurable, significant TCR clonotype sharing was found selectively between CD57^lo^PD-1^dim^ and CD57^lo^PD-1^hi^ as well as between CD57^hi^PD-1^hi^ and CD57^lo^PD-1^hi^ compartments. Interestingly no repertoire overlap was found between CD57^lo^PD-1^dim^ and CD57^hi^PD-1^hi^. Our data indicate a unidirectional transition from CD57^lo^PD-1^dim^ to CD57^lo^PD-1^hi^ to CD57^hi^PD-1^hi^. Importantly, CD57^hi^ TFH cells were characterized by i) increased expression of Bcl-6 and expression of genes associated with TFH development, ii) a shift from IL-21 to IL-4 production, which promotes the GC responses(11) and iii) an increased expression of genes encoding cytokines (IL-5 (46), IL-2 (47, 48)) involved in B cell differentiation to plasma cells. Therefore, the functional and spatial positioning profile suggests that CD57^hi^ TFH cells is the TFH subset providing B cell help in later steps of GC B cell development.

Differentiation of TFH cells was associated with dramatic changes in TCR expression. The absence of evidence for differential transcription of TCR subunits suggests that the downregulation of TCR in TFH cells is regulated post-translationally. Our imaging assay/analysis, which detects both surface and intracellular CD3, suggests that altered TCR recycling is not the reason for the observed TCR dynamics either. The presence of CD150 was associated with further downregulation of TCR. We have previously described the compromised *in vivo* division of CD150^lo^ compared to CD150^hi^ TFH cells in chronically SIV infected non-human primates (8). We assume that CD57^hi^ TFH cells, which mainly express a CD150^lo^ profile accompanied by significantly reduced expression of TCR and Ki67^hi^, have compromised capacity for *in vivo* division compared to CD57^lo^ TFH cells. Analysis of lymph node tissue sections and tissue derived cells from healthy individuals without history of cancer or infection revealed a similar to tonsillar TFH cell phenotypic profile, downregulation of TCR and relative localization of CD57^hi^ vs CD57^lo^ TFH cells. Our data imply that, at least, the described profiles is an intrinsic characteristic of TFH cells and not the consequence of local tissue inflammation.

Despite the downregulation of TCR, TFH cells express higher levels of integrins and CD151 suggesting that TFH cells are not characterized by a generalized reduction of IS components. Compared to CD57^lo^, CD57^hi^ TFH cells showed significantly i) smaller IS area, ii) lower numbers of TCR microclusters and synaptic TCR intensity (a measurement of synaptic TCR molecules) and iii) lower capacity for surrogate antigen uptake, a process defined as trogocytosis (49). Reduced trogocytosis, associated with reduced proliferation (50), could contribute to the compromised proliferative capacity found for CD57^hi^ TFH cells. We used total amount of tyrosine phosphorylation as a readout for TCR induced signaling at IS level. A lower TCR-induced synaptic tyrosine phosphorylation was found in CD57^hi^ TFH cells, a profile not associated with an overall compromised ability for tyrosine phosphorylation in CD57^hi^ compared to CD57^lo^ TFH cells (imaging analysis of pTyr). Supporting to this pattern is the increased potential for phosphatase activity, in CD57^hi^ compared to CD57^lo^ TFH cells. Compared to CD57^lo^, CD57^hi^ TFH cells express higher levels of SHP-2, an inhibitor of TCR signaling (51, 52), significantly lower levels of CD150, a receptor that can affect the phosphatase activity by recruiting either SHIP or SHP-2 phosphatases (33) and marginally higher levels of PTEN. Therefore, CD57^hi^ TFH cells have a potentially higher capacity for tyrosine dephosphorylation compared to CD57^lo^ TFH cells that could contribute to the observed pTyr profiles in peripheral synaptic microclusters, the sites where TCR signal is sustained (53).

Sequencing analysis, flow cytometry and biochemical assays revealed significantly increased proteasome activity in TFH compared to non/pre-TFH cells. TCR is ubiquitinylated before translocating into cSMAC (35) where TCR and TCR-elicited signals are sorted for degradation with proteasome activity been a critical force for this process (53). The limited free ubiquitin in TFH cells could lead to incompetent ubiquitination of TCR and IS proteins and their limited trafficking to cSMAC for degradation. Disruption of the proteasome activity in naïve cells resulted in more dispersed, peripheral type of IS, further supporting the role of proteasome in the type of IS by lymphoid organ CD4 T cells. The absence of an effect of MG132 in sorted TFH cells could be attributed to the inability of the inhibitor to control the very high proteasomal activity found in these cells or the fact that these cells may have already reached the threshold of peripheral clusters generation even in the absence of MG132.

IS-related protein degradation is found in the cSMAC area but not in the TCR microclusters (53). We hypothesize that the particular profile of the TFH IS is the outcome of a limited ubiquitination/translocation of TCR into cSMAC for degradation accompanied by higher local tyrosine phosphatase potential. Despite the significantly lower levels of TCR in TFH cells, particularly the CD57^hi^ ones, such mechanism could, at least in part, sustained TCR microclustering and TCR-induced signaling *in vivo*. In line with the concept that TCR signal quality and strength can affect the differentiation of effector cells(54), we propose that strong initial activation, proliferation of naïve and less differentiated cells is facilitated by a cSMAC type of IS (higher microcluster TCR and pTyr intensity)(55) while an enlarged pSMAC type of IS, mediated by upregulation of integrins and downregulation of TCR, supports the communication between CD57^hi^ TFH and GC-B cells by shaping the architecture of immunological synapse where GC B cells have also been found to form pSMAC oriented synapse for uptake of antigens (56) (Figure 8). Therefore, pSMAC TFH synapse is possibly a preferred architecture of IS within germinal center for the GC-TFH and B cell interaction.

**Figure 8:**
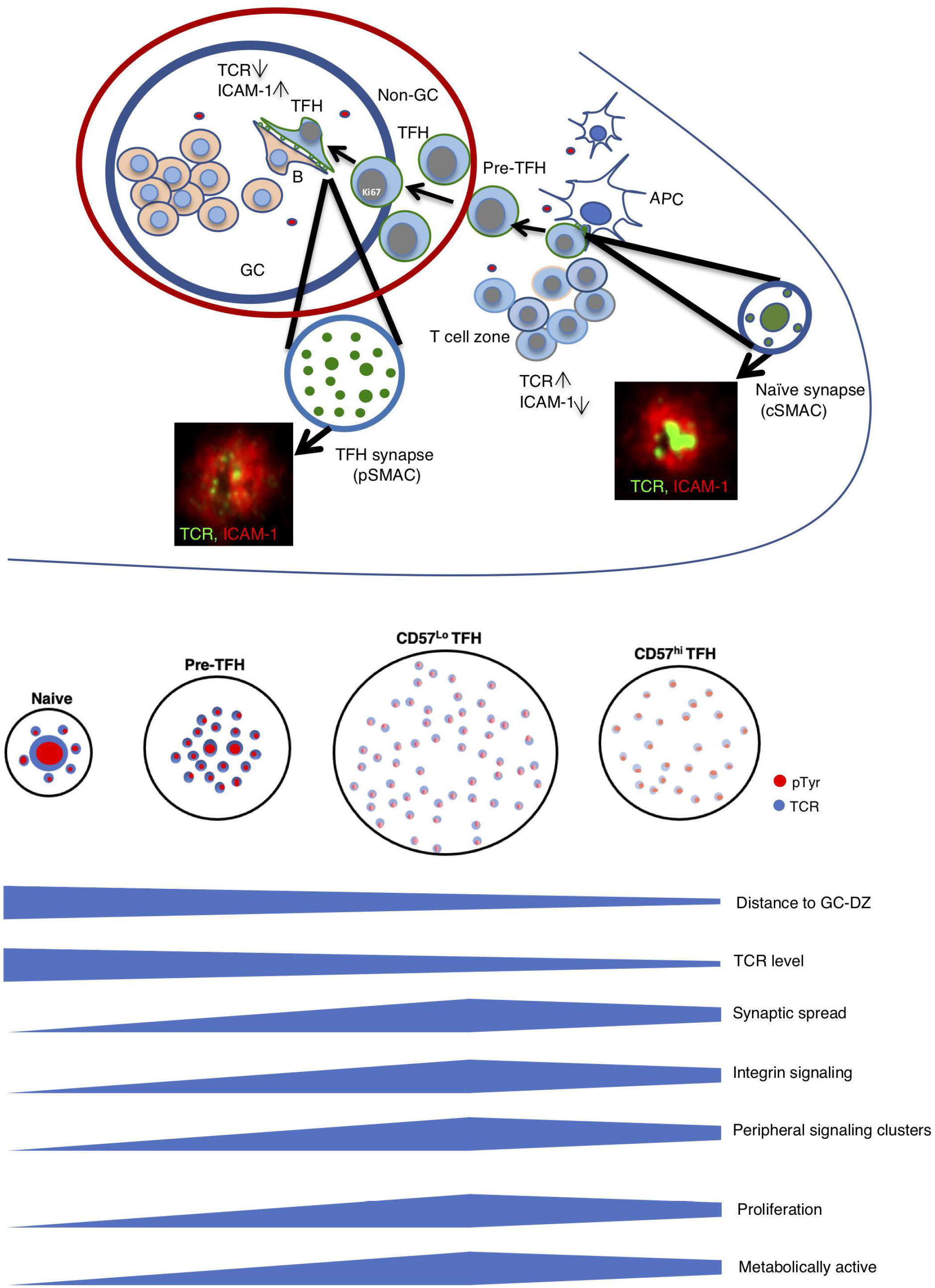
Differentiation towards TFH stage is characterized by dramatic changes in immunological synapse formation. The cartoon depicts the major changes observed at IS level during the transition from nai□ve to pre-TFH to CD57lo and CD57hi TFH cells. The relative location of these subsets to the follicular area is also shown. The majority of nai□ve cells form a cSMAC (represented by the centralized circle) type of IS with high intensity of synaptic TCR and pTyrhi proteins while the presence of cSMAC is becoming less evident with the CD4 T cell differentiation. The color intensity (red for pTyr and blue for TCR) reflects the relative intensity of these factors within the microclusters (small circles) and the size of the circle the relative size of the synaptic area in each cell subset. Horizontal bars represent the relative “size” of individual biological factors.

Whether the described distinct profile of CD57^hi^ TFH cells represents a stable, unique differentiation stage or a TFH subset that has recently interacted with antigenpresenting GC B cells and subsequently downregulated their TCRs and altered their transcriptional and phenotypic profile is not clear. Our data indicate that the described profiles apply to LN TFH cells too. Investigation of human LNs from different time points (i.e. using FNA preparations after vaccination (57)) or similar non-human primate TFH subsets in infected or vaccinated animals would be highly informative regarding this issue. The described TFH cell profiles could have important implications for our understanding of the role of TFH cells in HIV reservoir formation (42, 58) and the development of vaccines aiming to strength the humoral responses. Distinct localization of TFH subsets, presumably exposed to different microenvironment stimulation signals, is accompanied by differential expression of TCR levels and unique IS formation that could act as a modulator for the level of TFH activation. Furthermore, co-evolution of the metabolic program and TCR expression could promote a senescent status in particular TFH cells (i.e. CD57^hi^CD150^lo^) with a direct effect on the rate of latent virus reactivation. Alternatively, the lower expression of TCR could compromise the reactivation of latently infected cells by immunotherapies like bispecific antibodies aiming to eliminate infected TFH cells. Our data provide an immunological base for the characterization of “effective” TFH cells, potential targets of vaccine strategies, and help to define GC signatures that could inform for the spatial organization of specific cellular and molecular players for the development of immunogen-specific B cell responses.

## Methods

### Human material

Inguinal lymph nodes from 8 healthy subjects who underwent vascular (varicose vein stripping) and general (uncomplicated bilateral inguinal herniorrhaphy) surgery were used for cell suspensions preparation. 4 cervical and one mediastinal lymph nodes from healthy individuals were used for imaging of relevant cell populations. All lymph nodes were part of the surgical specimen obtained during elective vascular and cardiovascular surgery. Tonsils were obtained from anonymized discarded pathologic specimens from Children’s National Medical Center (CNMC) under the auspices of the Basic Science Core of the District of Columbia Developmental Center for AIDS Research. The CNMC Institutional Review Board determined that study of anonymized discarded tissues did not constitute ‘human subjects research.’

### Tissue Processing

Upon receipt, tonsils were washed with ice-cold medium R-10 (RPMI 1640 supplemented with 10% fetal bovine serum, 2 mM L-glutamine, 100 U/mL penicillin and 100 μg/mL streptomycin (Invitrogen), surrounding fatty tissue was removed and tissues were cut into small pieces. Part of tissue was immediately placed in fixative (10% formalin) and dedicated for paraffin embedded blocks (FFPE) followed by sectioning and analysis by multiplex confocal microscopy. Mononuclear cells were isolated by mechanical disruption followed by Ficoll-Paque density gradient centrifugation. Cells were then counted and frozen at 10×10^6^ cells/ml of freezing media (10% DMSO in fetal bovine serum) (Lonza). and stored in liquid nitrogen until further use.

### Antibodies

***Flow cytometry:*** directly conjugated antibodies were obtained from the following sources, *BD Biosciences:* CD57-FITC (NK-1), CD4-BV650 (SK3), CD45RO-BV786 (UCHL1), CD3-H7APC (SK7), OX40-BV650 (ACT35), CD19-APC (HIB19), CD45RO-BV786 (UCHL1), TCRαβ-APC (T10B9.1A-31), CD69-BV421 (FN50), CD4-BV650 (SK3), CXCR4-Cy5PE (12G5), CD151-BV786 (14A2.H1), CD18-BV421 (6.7), CD3-BB700 (SP34-2), Ki67-BV421 (B56), and Bcl-6-PE (K112-91), *Biolegend:* CD27-BV605 (O323), PD-1-BV711 (EH12.2H7), ICOS-Pacific Blue (C398.4A), CD19-BV570 (HIB19), CD27-BV605 (O323), PD-1 BV711 (EH12.2H7), CD150-PE (A12/7D4), CD20-BV570 (2H7), and CD20-BV570 (2H7), *eBioscience:* TIGIT-APC (MBSA43) and CXCR5-Cy7PE (MU5UBEE), c-Maf eF660 (sym0F1), and TIGIT-APC (MBSA43), *Abcam:* K-48 linkage specific ubiquitin antibody Alexa-488 (EP8589), *Cell Signaling:* PTEN-Alx488 (rabbit polyclonal, #9560).

***Cytof:*** metal isotopes conjugated antibodies were obtained from *Biolegend:* CD8-113In (RPA-Y8), ICOS-143Nd (C398.4A), CD3ε-149Sm (SK7), CXCR3-154Sm (GO25H7), CD25-158Gd (M-A251), CCR7-159Tb (GO43H7), CD150-167Er (A12), TCRγδ-173Yb (B1), *BD:* CD151-142Nd (14A2.h1), CD3αβ-166Er (T10B9.1A-31), CD57-194Pt (G10F5), *DVS:* CD45-141Pr (HI30), Cd38-144Nd (HIT2), IgD-146Nd (IA6-2), Ox40150nd (ACT35), TIGIT-153Eu (MBSA43), CD27-155Gd (L128), CXCR4-156gd (12G5), Tbet-160Gd (4B10), Ki67-161Dy (B56), CD69-162Dy (FN50), Bcl-6-163Dy (K112-91), CD45Ro-165HO (UCHL-1), CD45RA-170Er (HI100), CD40L-168Er (24–31), CCR5-171Yb (NP-6G4), HLA-DR-174Yb (L243), PD-1-175Lu (EH12.2H7), CD4-176Yb (RPA-T4).

***Scanning confocal microscopy:*** *BD Biosciences*: Ki-67-BV421 (B56), CD57-Alexa680 (clone NK-1), CD20-ef615 (L26), *R&D Systems:* PD-1 (FAB7115G)); CD4-Alexa488 (FAB8165G), *Dako:* CD3 (F7.2.38), CD68 (KP1). *Cell Signaling:* Phospho-Tyrosine (9414S), *Sigma:* FDC (CNA.42). *Invitrogen:* Anti-mouse IgG-Alexa647 conjugated secondary antibody (A21463), goat anti-mouse IgG1-Alexa594 conjugated secondary antibody (A21125). *Abcam:* IgD (EPR6146).

***TIRF:*** *eBioscience*: CD3 (OKT3), *Millipore:* LBPA (6C4), *Cell Signaling:* Phospho-Tyrosine (9414S).

***Western blotting:*** *Covance*: mono-ubiquitination specific antibody (3–39). *Thermo Fisher Scientific:* β-actin (BA3R), *SouthernBiotech:* HRP-conjugated anti-mouse secondary antibody (1010-05), *Boster:* anti-K48-linkage specific ubiquitin antibody (IBH-21).

### Chemicals and reagents

Streptavidin-(Jackson ImmunoResearch, 016-000-084), JOJO-1 iodide (Life Technologies), Fluoromont-G (0100-01 Southern Biotech), Aqua fluorescence dye- (Invitrogen L34966), DOPC (#850475C) and DGS-NTA (#790404C) (Nickel salt), lipids (Avanti Polar Lipids), Silica beads (SS06N, Bangs Lab), NHS-ester form of Alexa 488 and 647 dyes (Thermo fisher Scientific), λ-phosphatase (NEB), Phalloidin (Invitrogen, A22287), Aqua Dead Cell Stain (Thermo Fisher, Catalog L34965), 2-NBDG (N13195-Thermo Fisher Scientific), Phalloidin-Alexa647 (A22287 Thermo), mitotracker green (M7514, Thermo Fisher Scientific), Fluorogenic Substrate III (Calbiochem, 5391425MG), MG132 (Calbiochem, 474790-1MG), Lactacystin (Fisher Scientific, APT280). Purification of histidine tagged ICAM-1 and has been described before (59). HBS BSA buffer was prepared according to the recipe published by Dustin ml. Basically, it contains 20 mM HEPES, pH 7.2, 137 mM NaCl, 5 mM KCl, 0.7 mM Na_2_HPO_4_, 6 mM d-glucose, 2 mM MgCl_2_, 1 mM CaCl_2_, 1% (w/v) BSA. Etoposide was from Sigma (E1383-25MG).

### Transcriptome analysis

***Microarray Analysis:*** reverse transcription reactions were performed to obtain cDNAs which were hybridized to the Illumina Human HT-12 version 4 Expression BeadChip according to the manufacturer’s instruction and quantified using an Illumina iScan System. The data were collected with Illumina GenomeStudio software (illumine.com). Analysis of the genome array output data was conducted using the R statistical language and the Bioconductor suite. First, arrays displaying unusually low median intensity, low variability, or low correlation relative to the bulk of the arrays were tagged as outliers and were discarded from the rest of the analysis. Quantile normalization followed by a log2 transformation using the Bioconductor package LIMMA was applied to raw microarrays intensities. The LIMMA package was used to fit a linear model to each probe and to perform a (moderated) Student’s *t*-test to assess the association of gene-expression to a phenotype of interest. For data mining and functional analyses, genes that satisfied a p-value <0.05 were selected. Probes that do not map to annotated RefSeq genes and control probes were removed. When indicated, the proportions of false positives were controlled using the Benjamini and Hochberg method. ***Pathway analysis:*** gene Set Enrichment Analysis (GSEA) using the MSigDB signatures (http://software.broadinstitute.org/gsea/msigdb/index.jsp) was performed to identify pathways associated with a sorted Tfh cell compared to another. GSEA is a statistical method to determine whether members of a particular gene set preferentially occur toward the top or bottom of a ranked-ordered gene list where genes are ranked by the strength of their association with the outcome of interest. More specifically, GSEA calculates an enrichment score (NES) that reflects the degree to which a set of genes is overrepresented among genes differentially expressed. The significance of an observed NES is obtained by permutation testing: resorting the gene list to determine how often an observed NES occurs by chance. Leading Edge analysis is performed to examine the particular genes of a gene-set contributing the most to the enrichment. ***Network Mapping/Visualization:*** GeneMania Networks (Genemania.org) were plotted to represent co-expression of enriched genes from pathways of interest. Gene Mania is a flexible, user-friendly web interface for generating hypotheses about gene function, analyzing gene lists and prioritizing genes for functional assays. Given a query list, GeneMANIA extends the list with functionally similar genes that it identifies using available genomics and proteomics data. GeneMANIA also reports weights that indicate the predictive value of each selected data set for the query. The Cytoscape (cytoscape.org) plugin was used to plot the networks. ***Single cell TCR clonotype analysis:*** sorted total memory (CD27^hi/lo^CD45RO^hi^) tonsillar CD4 T cells (10,000 targeted single cells per sample) were used to generate sequencing libraries for α/β TCR repertoire and 5’ transcriptome analysis. Library preparation was performed according to the manufacturer’s specification and the libraries were subsequently sequenced on the Illumina NovaSeq at 2×150 base pairs on an S1 flow cell. Characterization of single cell transcriptomes and identification of α/β TCR clonotypes for each cell was analyzed with Cellranger software. Cells for which a transcriptome and α/β TCR were recovered where selected for clonotype analysis, yielding 1022, 2930, 1609, 2710, 2653 and 3807 cells, respectively. Clonal overlap between each subset on a per-sample basis was measured by the Morisita-Horn similarity index, which incorporates clonotype abundance and hierarchy.

### Polychromatic Flow cytometry

***Phenotypic analysis:*** 2-3×10^6^ cells were thawed out and rested for 2h in a cell culture incubator before further use. Fresh tonsil-derived cells were also used for some experiments. Cells were washed with PBS, BSA (1%), incubated (5min, RT) with a viability dye (Aqua-dye, Invitrogen) and surface stained with titrated amounts of antibodies against i) ICOS-PB, CD19-BV570, CD27-BV605, OX40 BV650, PD-1 BV711, CD45RO BV786, CD150-PE, CD4 Cy55PE, TCR ab-APC, CD3 epsilon-H7APC and CD57 Alexa Fluor 680, ii) CD57-FITC, CD69-BV421, CD20-BV570, CD27-BV605, CD4-BV650, PD-1 BV711, CD45RO ECD, CXCR4 Cy5PE, CD151-BV786, CD18 BV421, CXCR5 Cy7PE, TIGIT-APC and CD3 H7APC. Following a washing step, cells were fixed with 1% paraformaldehyde. For the detection of PTEN, following surface staining with Aqua, and antibodies against CD3, CD4, CD27, CD45RO, PD-1, CD57 cells were washed and fixed/permeabilized with BD buffer II (cat. Number 558052) and stained intracellularly with anti-PTEN for 30 min. A similar protocol was performed for the detection of p53 after stimulating cells for 6h with 50 or 200 uM etoposide. Active form of LFA-1 was measured by looking at binding of soluble His-ICAM-1 for 30 min at 37°C in HBS BSA containing Ca^++^ and Mg^++^. ***Transcription factors (ICS) analysis:*** transcription factor staining was performed using the Transcription Factor buffer set (BD Pharmingen) per manufacturer’s instructions. Briefly, 2 million tonsil-derived lymphocytes were thawed in R10 and rested for 2hr at 37°C before being stained with the LIVE/DEAD fixable Aqua Dead Cell Stain and the following cocktail of surface antibodies: CD3-BB700, CD20-BV570, CD27-BV605, CD4-BV650, PD-1 BV711, CD45RO BV786, CD57 AF594, and CXCR5 Cy7PE. After a 20-minute incubation, cells were washed, permeabilized per BD protocol and stained intracellularly using the following antibodies: TOX2-FITC, Ki67 BV421, Bcl-6 PE and cMaf eF660. For ubiquitin staining, the Alexa-488 conjugated K-48 linkage specific ubiquitin antibody was used. Phalloidin-Alexa647 used to detect actin in some ICS experiments. Following a washing step, cells were fixed with 1% paraformaldehyde. ***Mitochondria and glucose uptake analysis:*** TMRE staining was performed as previously described (24)). Briefly, 2 million tonsil-derived lymphocytes were thawed in R10 (RPMI+10% serum) and rested for 2hr at 37°C before being stained with the LIVE/DEAD fixable Aqua Dead Cell Stain and the following cocktail of surface antibodies: antibodies CD57-FITC, CD27 BV605, CD4 BV650, PD-1 BV711, CD45RO BV786, CD19 APC and CD3 H7APC. Cells were surface stained for 20 minutes, washed, re-suspended in RPMI containing 5uM of either TMRE dye (Invitrogen, T669) or FFCP (Sigma-Aldrich) and incubated for 30 further minutes. For Mitotraker Green FM analysis, after surface staining with the same antibodies panel, cells were incubated in serum free RPMI medium containing 25nM Mitotracker green for 30 min. To measure glucose uptake, cells were incubated with 50 μM 2-NBDG (Thermo Fisher Scientific) in glucose free RPMI medium at 37C for 30 min after applying the above surface panel staining. After staining with the relevant chemicals, cells were washed and acquired without fixation on a BD LSR Fortessa X-50 flow cytometer. ***Data acquisition:*** 0.5-1×10^6^ events were collected in each case on a BD LSR Fortessa X-50 flow cytometer (BD Immunocytometry Systems). Electronic compensation was performed with antibody capture beads (BD Biosciences). Data were analyzed using FlowJo Version 9.9.4 (Tree Star). Forward scatter area vs. forward scatter height was used to gate out cell aggregates.

### Mass Cytometry (CyTOF)

Cryopreserved lymph node mononuclear cells (LNMCs) were thawed and resuspended in complete RPMI medium (Gibco; Life Technologies; 10% heat inactivated FBS [Institut de Biotechnologies Jacques Boy], 100 IU/ml penicillin, and 100 μg/ml streptomycin [BioConcept]). Cells were washed twice and then the viability of cells in 500 μl of PBS was assessed by incubation with 50 μM cisplatin (Pt-198; Fluidigm) for 5 min at RT and quenched with 500 μl fetal bovine serum. Next, cells were incubated for 30 min at 4 °C with a 50 μl cocktail of cell surface metal conjugated antibodies (Fluidigm). Cells were washed and fixed for 10 min at RT with 2.4% PFA. Next, cells were permeabilized for 45 min at 4°C with Foxp3 Fixation/Permeabilization kit (eBioscience), washed and stained at 4°C for 30 min with a 50 μl cocktail of transcription factor metal conjugated antibodies. Cells were washed and fixed for 10 min at RT with 2.4% PFA. Total cells were identified by DNA intercalation (1 μM Cell-ID Intercalator, Fluidigm) in 2% PFA at 4°C overnight. Labeled samples were assessed by the Helios mass cytometer instrument (Fluidigm) using a flow rate of 0.045 ml/min. Analysis of the mass cytometry data was performed on FCS files that were normalized to the EQ Four Element Calibration Beads using the CyTOF software. For conventional cytometric analysis of CD4 T cell populations, FCS files were imported into Cytobank Data Analysis Software or FlowJo v10.4.2 (Treestar, Inc., Ashland, CR). Unsupervised clustering was conducted using FlowSOM (BuiltSOM function in FlowSOM package) and t-SNE analysis using Cytobank and R software packages.

### Cell sorting

***TIRF microscopy:*** frozen cells (50×10^6^) from tonsil were thawed, rested (2h) and stained for aqua, CD20, CD4, CXCR5 and ICOS. Live CD4 T cells were sorted for ICOS-1^lo^CXCR5^lo^, ICOS-1^dim^CXCR5^dim^) and ICOS^high^CXCR5^high^. For some experiments, ICOS^high^CXCR5^high^ cells were further sorted for CD57^low^ and CD57^high^ subsets. For comparison, we also sorted TFH cells based on PD1 and CXCR5 expression in a control experiment and found out that PD-1 and ICOS represent same populations of TFH in terms of phenotype and synapse formation. In general, we used ICOS instead of PD-1 to exclude any interference from PD-1 mediated inhibitory effect. Likewise, we avoided use of anti-CD3 antibodies in the sorting to avoid TCR stimulation. We sorted 1-2 million cells from each cell types for experiments involving FCS2 chamber (Bioptechs) and 100,000-200,00 cells for experiments involving 8-well chamber (Lab-Tek). Sorted cells were rested for 1hr at 37°C before the start of experiments. ***Gene sequencing:*** memory CD4 T cell subsets were sorted based on the expression of PD-1 and CD57 markers. Again, we avoided using anti-CD3 antibodies in the sorting to avoid TCR stimulation. 50,000 to 100,000 sorted cells were used for downstream sequencing analysis

### Luminex-secreted cytokines

The levels (pg/ml) of secreted cytokines after *in vitro* stimulation of sorted CD4 T cell subsets with PMA (10ng/ml)/ionomycin (1ug/ml) for 14-16h were measured using Luminex technology according to the manufacturer’s instructions (customized kit from R&D).

### Preparation of glass supported lipid bilayer (SLB)

Liposomes containing 6.25% DGS-NTA and 2% Cap-Biotin-DOPC lipids were prepared using an extruder (Avanti Polar Lipids, Inc.) according to manufacturer’s instructions. Glass-supported lipid bilayer was prepared in FCS2 chamber (Bioptechs) as described previously (29, 59). For some experiments supported lipid bilayer was formed in 8-well glass bottom chamber (Lab-Tek) as described previously. Briefly, liposomes were applied to charged-cover glasses (erased with 30% H_2_O_2_ and 70% H_2_SO_4_) for about 30 seconds to form planar lipid bilayer. The lipid bilayer was washed with HBS BSA buffer (Dustin ml) before incorporating the ligands. Then, streptavidin (5 g/ml) was incorporated into the bilayer (30min/RT) followed by Alexa dye conjugated monobiotinylated anti-CD3 antibodies (2.5μg/ml), histidine tagged CD80 (100 molecules/ μm^2^), and ICAM-1 (100 molecules/ μm^2^). After 30 min incubation with ligands, the bilayer was washed and blocked with 5% casein for 20 min. The chamber with lipid bilayer is now ready for imaging in TIRF microscope.

### Supported lipid bilayer on silica beads

Lipid bilayers were formed on 5μm sized silica beads (Bangs Laboratories, SS06N) as described in (60). Liposomes containing Cap-biotin and NTA lipids were mixed by vertexing with beads at 2:1 ratio (by volume). Bilayer coated beads were washed with HBS buffer by spinning at 600g for 2 min. After washing, the beads were incubated with Streptavidin (30 min RT) before adding mono-biotinylated OKT3, CD80 and ICAM-1 for another 30 min at RT in HBS BSA buffer containing Ca^++^ and Mg^++^. The ligand coated beads are now ready for cell stimulation. Sorted T cells were stimulated with ligand-coated beads in 1:2 ratio of cells to beads. For short-term stimulation (up to 2hr) the cells were incubated in HBS BSA buffer with Ca^2+^ and Mg^2+^ at 37°C without CO_2_.

### LBPA staining

Sorted cells were made to interact with the ligand coated lipid bilayer for 15 min for stable synapse formation. The cells were then fixed with 2% PFA for 20 min at RT. This was followed by incubation with 50mM NH_4_Cl for 10 min. Then, cells were permeabilized with 0.05% Saponin and stained with anti-LBPA antibody (primary) followed by Alexa594 conjugated anti-mouse IgG secondary antibody.

### λ-phosphatase treatment

To test the specificity of phospho-tyrosine antibody, λ-phosphatase use used as per manufacturer’s instruction (NEB). FFPE tonsil section was processed for antigen retrieval followed by staining with phospho-tyrosine antibody in the presence or absence of λ-phosphatase (50 units/ml) in Mn^2+^ containing buffer. After 1hr incubation at 37°C, the tissue was washed with PBS and stained with CD20 antibody for confocal imaging.

### MG132 treatment

Treatment of tonsillar T cells with MG132 was performed as described previously (35). Briefly, sorted T cells were administered for 2hr at 37°C at 10μM MG132 in culture media to deplete cells free of ubiquitin. They were then washed and resuspended with HBS BSA buffer and incubated on glass supported bilayer for TIRF imaging in presence of the inhibitor. Control cells were treated with DMSO.

### Antigen uptake assay

Sorted cells were rested for 1hr at 37°C and then incubated with beads containing anti-CD3-Alexa 488 (2.5 μg/ml), CD80 and ICAM-1. After 1hr incubation at 37°C, live cells were stained for surface panel as described above and then analyzed for the uptake of α-CD3-Alexa488 by flow cytometry.

### Western blotting

Detection of mono/poly-ubiquitination was performed by western blotting by using protocol as described earlier (59). Briefly, 2 million sorted tonsillar cells were lysed in 1X SDS containing Laemmli buffer. Cell lysate was loaded into 4-15% precast gels (Bio-Rad) for electrophoretic separation. Proteins were transferred onto PVDF membrane (Invitrogen) and blotted for ubiquitin and actin.

### Proteasome activity assay

Sorted tonsillar cells (~2 × 10^6^) were resuspended in 100 μl of lysis buffer (13mM Tris-HCl and 5mM MgCl_2_, pH 7.8) and subjected to three rounds of freeze-thaw lysis. Cell lysate was collected after spinning at 13000 rpm for 10 min at 4°C and protein concentration was determined by Nanodrop. The cell lysate was brought up to a final volume of 200 μl by adding lysis buffer supplemented with 5mM ATP, 0.5mM DYY, 5mM EDTA and 100 μM final concentration of fluorogenic substrate III. Fluorescence emission at 460nm (excitation 355nm) was recorded by ELISA plate reader at 1hr postincubation at 37°C. Amount of AMC released (in nanomoles) was measured using a standard curve generated from free AMC (Calbiochem). The proteasomal activity was then normalized to protein concentration. Reaction with a standard kit in the presence or absence of proteasomal inhibitor 10 μM Lactacystin was used for positive and negative control.

### Microscopy studies

***Confocal Microscopy:*** tissue blocks tissue sections were subjected to deparaffinization followed by antigen retrieval in Borg Decloaker RTU (Biocare Medical) at 110°C for 15 minutes. Tissue sections were then treated for permeabilization (0.3% Tritox-100 in PBS) followed by staining with primary antibodies (O/N at 4°C), secondary antibodies (2hr at RT), blocking with 10% goat serum (1hr at RT) and then conjugated antibodies (2hr at RT). Finally, slides were then stained for 20 min with JOJO-1 (Life Technologies) and mounted with with Fluoromount-G (Southern Biotech). Stained slides were imaged on a NIKON (C2 or A1 systems) inverted confocal microscope equipped with 40X, 1.3 NA oil objective lens. Image acquisition was performed with NIS-elements software and analyzed in Imaris software version 8.2 (Bitplane). Spectral spillover between channels was corrected through live spectral unmixing in NIS using data acquired from samples stained with single fluorochromes.

### Microscopy studies

***Confocal Microscopy:*** imaging of FFPE tissue sections was performed as described previously (61). Histocytometry analysis was performed as published earlier (62, 63). Briefly, imaging datasets were segmented post-acquisition based on nuclear staining and average voxel intensities for all channels were extrapolated in Imaris. Channel statistics were exported to csv (comma separated values) files format and analyzed in Flow Jo version 10. ***Cell-cell Distance analysis:*** imaged tissue sections were used in a comparative distance analysis between specific cell populations and structural regions. CD20, Ki67, and FDC were used to manually draw and generate surfaces demarcating the regions of the follicle, the germinal center, and the follicular dark zone and light zone for follicular areas selected for having well-defined germinal centers and well-defined and separated light zone/dark zone morphology. Making use of the ImarisXT license, a Python script was used on each image to generate spots whose center points matched the X/Y coordinates of the nuclear center of each cell in several populations determined via histocytometry performed on each image. For each selected follicle, cells were further filtered and placed in subgroupings based on whether their centers lie inside those regions. In Imaris, for each follicle the module XTDistanceTransform was used to determine the shortest distance in μm of each cell center in the CD57^hi^ and CD57^-^ GC TFH populations to various features. The shortest distance to any part of the dark zone region or FDC network for each cell was calculated. Cells whose centers lie within or on the border of the dark zone are demarcated as having a distance of zero μm to the dark zone. Also determined was the shortest distance to any part of the non-GC region of the follicle, which was determined by measuring each cell’s distance to the border of the GC. Cell centers located outside the GC or on the border of the GC are likewise marked with a distance value of zero μm. The shortest distance of each GC TFH to the center coordinates of any cell belonging to the Light Zone CD20^hi^Ki67^dim^ or CD20^hi^Ki67^hi^ groupings was calculated. In the tonsils, additionally, the shortest distance was determined for each GC TFH to the center of any Light Zone CD68^hi^ cell as well as the distance between PD-1^hi^CD57^lo^ki67^hi^ and PD-1^hi^CD57^lo^Ki67^lo^ to FDC. The distance values of each comparison listed above was recorded separately for the CD57^lo^ and CD57^hi^ GC TFH populations and exported through the Statistics tab in Imaris.

***Volumetric 3D imaging:*** multiplex volumetric tissue staining was performed using protocol published previously (64). Briefly, Tonsil sections (1-5 mm cubes) were fixed in 1% PFA at 4°C O/N. Fixed tissues were permeabilized in 0.3% Triton X-100 containing PBS for 24 hours at 37°C. Next, conjugated CD20, CD4, Ki67, PD-1 and CD57 were used for deep tissue staining by incubating at 37°C in orbital shaker for 3-5 days. Stained tissues were washed in PBS+0.2% Triton X-100 and then subjected to tissue clearing in Histodenz containing clearing solution for 3 days at 37°C. Cleared tissues were mounted with the clearing solution in 8-well glass bottom chambers (Lab-Tek) and imaged with 20X water objective lens in a Nikon A1 confocal microscope. ***TIRF microscopy:*** FCS2/8-well chamber containing the lipid bilayer was brought to 37°C before injecting the cells. Sorted cells (after 1hr rest) were made to interact with the ligand-coated bilayer for up to 1hr. Cell stimulation was carried out in HBS BSA buffer containing Ca^2+^ and Mg^2+^. Formation of immunological synapse was captured in live cell imaging by automated TIRF set up designed in Nikon C2si microscope equipped with 100X 1.45NA oil objective lens and ANDOIR camera. Adhesion of T cells to the lipid bilayer was determined by the dark interference image in surface reflection interference contrast (SRIC) microscopy. To perform intracellular phospho-protein staining, the lipid bilayer with interacting T cells were fixed with 2% PFA and then permeabilized with 0.05% Saponin followed by antibody staining. Quantification of the number and intensity of TCR, pTyr and LBPA cluster at the synapse was done using the software Icy. Quantification of spread area of synapse was done by measuring surface reflection interference contrast (SRIC) images in Image J and Metamorph software after background subtraction. Statistical analysis was performed in PRISM. For recording the movies for synapse formation, the live images of interacting cells were captured every 1 sec up to 10 min time.

### Statistics

Heatmap (Figure 1G) was created using ggplot2 R package. Graphs were prepared using Prism 7.0d. Flow-cytometry and histocytometry generated data were analyzed using the Wilcoxon test. For TIRF generated data, statistical significance was assessed using a proper test given data normality (Shapiro-Wilk test and QQ plots) and equality of variance (Levene’s test) among the different populations. ANOVA for imbalanced data (if normality and equivalence of variance hold) and Welch’s ANOVA (when equivalence of variance was violated) was applied. To identify pairs for which, the hypothesis of equal means does not hold, pairwise t-test was applied. Benjamini-Hochberg correction was applied for multiple comparisons. Hence, adjusted p-values are provided. If both conditions were violated, Kruskal-Wallis test was applied. Conover-Inman test was used to find the cell population pairs that do not originate from the same distribution whenever Kruskal-Wallis test hypothesis is rejected. Again, the same correction is enforced for the multiple comparisons accordingly. Results were considered to be significant for (adjusted) p-value < 0.05. Statistical analysis was performed using the R statistical package.

### Study approval

All protocols involving human samples (lymph nodes) were reviewed and approved by the Institutional Review Boards at the relevant institutions (Centre Hospitalier Universitaire Vaudois-CHUV, Lausanne, Switzerland and the Medical School, University of Thessaly, Greece). Participant IDs were randomized, and the samples were randomly numbered to perform the experiments. Signed informed consent was obtained from all participants in accordance with the Declaration of Helsinki.

## Acknowledgements

The authors would like to thank Dr Roederer, Margaret Beddal (Immunotechnology Section VRC) and Megha Kamath (cellular Section VRC) for their help with antibody conjugations and Luminex assay. This research was supported by the Intramural Research Program of the Vaccine Research Center, NIAID, National Institutes of Health.

## Author contributions

KP, EM, AC, CF, KG, AX, and GF performed the experiments, did the analysis and reviewed the manuscript. VP performed the statistical analysis. AP, MI and GP provided tissue and cell material and reviewed/edited the manuscript. RP, GP and RAK provided critical help for the interpretation of the results and reviewed/edited the manuscript. KP, EM, CF, KG and CP wrote the manuscript. CP conceived the study and designed the experiments.

## Data and materials availability

the sequencing data discussed in the manuscript have been deposited (GSE145418).

