## Supplementary figure legends for "Human follicular CD4 T cell function is defined by specific molecular, positional and TCR dynamic signatures"

**Supplemental Figure 1. Imaging analysis of tonsillar follicular cell subsets. (A)** Single imaged planes generated during the clear 3D imaging of three tonsillar follicular areas (ki67-magenta, PD-1-green and CD57-red). The corresponding supplementary video is also indicated under the images. **(B)** The computational workflow used for the distance analysis of relevant cell populations. **(C)** Graph showing the mean shortest distance of $CD57^{lo}$ and $CD57^{hi}$ TFH cells to non-GC follicular region in 20 follicles from four tonsils. Representative histograms showing the distribution (number of cells) of $CD57^{lo}$ and $CD57^{hi}$ TFH cells with respect to their distance from the non-GC follicular region in one follicle. **(D)** Representative confocal image showing the computationally generated FDC network surface (in green), and Ki67 expression in a follicular area, used for distance analysis (upper panel) and graph showing the mean shortest distance of $CD57^{lo}$ and $CD57^{hi}$ TFH cells to FDC network in 20 follicles from four tonsils (lower panel).

**Supplemental Figure 2. Phenotypic and transcriptional analysis of tonsillar follicular CD4 T cell subsets. (A)** Flow cytometry plots showing the expression of several surface receptors or transcription factors in tonsillar naïve and memory CD4 T cell subsets. **(B)** Correlation (linear regression) between PD-$1^{hi}$$CD57^{lo}$ and other CD4 T cell tonsillar subsets (n=20 tonsils analyzed). **(C)** Plots showing the relative frequency of tonsillar CD4 T cell subsets from tonsils (n=10) characterized by high expression of individual surface receptors/ (**D)** Expression of surface receptors per cell (judged by Mean Fluorescence Intensity, MFI) in tonsillar CD4 T cell subsets expressing a "$receptor^{hi}$" phenotype (n=10 tonsils). **(E)** Plots showing the relative frequency of tonsillar CD4 T cell subsets characterized by high expression of nuclear factors (n=10 tonsils analyzed). **(F)** Flow cytometry plot showing the CD4 T cell subsets used for sequencing analysis (left panel) and the PCA analysis for the four indicated populations (right panel). **(G)** Heatmaps representing the top 50 differentially expressed genes ($p<0.05$) between $CD57^{lo}PD1^{hi}$ and $CD57^{lo}PD1^{dim}$, $CD57^{lo}PD1^{dim}$ and $CD57^{hi}PD1^{hi}$, and $CD57^{lo}PD1^{hi}$ and $CD57^{hi}PD1^{hi}$ (left to right). Rows represent genes and columns represent samples. Hierarchical clustering (Euclidian distance, complete linkage) was used to regroup samples with similar expression. Gene expression is represented as a gene-wise standardized expression (Z-score). Red and blue correspond to up- and down-regulated genes respectively. Color annotation on top of the heatmap represents sorted population.

**Supplemental Figure 3. Functional analysis of tonsillar CD4 T cell subsets. (A)** Gene Set Enrichment Analysis (GSEA) was used to test the enrichment of pathways between $CD57^{lo}PD1^{hi}$ and $CD57^{lo}PD1^{dim}$ (upper panel) and $CD57^{lo}PD1^{hi}$ and $CD57^{hi}PD1^{hi}$ (lower panel) sorted populations (p-val<0.05). Bar-plot represents positively enriched (red, up in $CD57^{lo}PD1^{hi}$) and negatively enriched (blue, down in $CD57^{lo}PD1^{hi}$) pathways. Pathways are plotted on the y-axis and the NES score on the x-axis. **(B)** GSEA was used to assess the enrichment of TFH specific pathways between $CD57^{hi}PD1^{hi}$ and $CD57^{lo}PD1^{hi}$ sorted populations. Pathway Heatmap illustrating the normalized enrichment score (NES) of the top 12 TFH genesets (GSEA p-value $\leq 0.05$; MSigDB: c7) between $CD57^{hi}PD1^{hi}$ and $CD57^{lo}PD1^{hi}$ sorted populations. Red and blue squares represent positive or negative enrichment of a pathway among the genes up-regulated and down-regulated respectively between the 2 groups. **(C)** A co-expression network of the leading-edge genes of TFH pathways was plotted to highlight their interaction. Circular nodes represent genes and edges reflect the association between these features. Red nodes indicate these genes are upregulated in $CD57^{hi}PD1^{hi}$ compared to $CD57^{lo}PD1^{hi}$. **(D)** Flow cytometry showing the expression of TMRE (upper left panel) and Mitotracker GreenFM (upper middle panel) in tonsillar naïve (solid green line) and $CD57^{hi}PD\text{-}1^{hi}$ TFH cells (solid red line). The corresponding FMO levels are presented in solid (naïve) and dotted ($CD57^{hi}PD\text{-}1^{hi}$ TFH cells) black lines, for comparison. The levels of TMRE in cells treated with FCCP, a reagent that collapses the mitochondrial membrane potential, are also shown for naïve (dotted green line) and $CD57^{hi}PD\text{-}1^{hi}$ TFH cells (dotted red line). Histogram showing the level of NBDG (right panel) uptake by naïve (solid green line) and $CD57^{hi}PD\text{-}1^{hi}$ TFH cells (solid red line). The corresponding FMO measurements are also shown. **(E)** The expression of c-myc (MFI) in tonsillar CD4 T cell subsets, marked with different symbols, is shown (n=5 tonsils analyzed). **(F)** Expression of p53 in CD4 T cell subsets from two tonsils in the presence of etoposide for 3 or 6h. **(G)** Gene profiles generated from the array analysis of sorted bulk tonsillar CD4 T cell subsets were used for the categorization of single cells analyzed by 10x platform. A representative UMAP analysis showing the distribution of the identified CD4 T cell subsets (upper panel). The frequencies of the individual subsets are shown in the lower panel. **(H)** The levels of secreted cytokines after *in vitro* stimulation of sorted tonsillar CD4 subsets with PMA/ionomycin are shown.

**Supplemental Figure 4. Analysis of CD3 dynamics in tonsillar CD4 T cell subsets. (A)** Flow cytometry 2D plots showing the FMO staining for the CD3ε and CD3αβ chains (upper panel) and the expression of CD3 chains in total live cells and with respect to the expression of PD-1 or CD57 (lower panel). **(B)** Accumulated data showing the levels of CD3ε and CD3αβ in tonsillar CD4 T cell subsets marked with different symbols (second cohort of tonsils analyzed, n=6). **(C)** Flow cytometry plots showing the expression of CD150 in naïve and memory CD4 T cell subsets (red rectangles show the gating for the identification of $CD150^{hi}$ cells). **(D)** Accumulated data showing the relative frequency of $CD150^{hi}$ cells in tonsillar CD4 T cell subsets marked with different symbols (left panel) and the expression of CD3ε and CD3αβ chains (MFI) in $CD150^{hi\ or\ lo}$ TFH cells (right panel) is shown (second cohort of tonsils analyzed, n=6). The Wilcoxon test used for data analysis. **(E)** GSEA was used to assess the enrichment of TCR signaling pathways (PID c2, MSigDB) between $CD57^{lo}PD1^{hi}$ and $CD57^{lo}PD1^{dim}$ sorted populations. A co-expression network of the leading-edge genes of TCR signaling was plotted to highlight their interaction. Gene Mania algorithm was used to infer network connections and co-expression. Circular nodes represent genes and edges reflect the association between these features. Color of the edge highlights the association between nodes. All genes are down regulated in $CD57^{lo}PD1^{hi}$ compared to $CD57^{lo}PD1^{dim}$. Biological annotation (diamond nodes) reflects function of genes.

**Supplemental Figure 5. Analysis of human lymph node CD4 T cell subsets. (A)** Cytof plots showing the gating for the identification of relevant human LN CD4 T cell subsets. **(B)** Confocal images (scale bar: 100 um) showing the localization of $IgD^{hi}$ (cyan), $FDC^{hi}$ (white), $Ki67^{hi}$ (magenta), $CD20^{hi/dim}$ (blue), $PD\text{-}1^{hi}$ (red) and $CD57^{hi}$ (green) cells in a follicular area from healthy LN. Solid lines depict the borders for particular follicular areas (white: Non-GC, yellow: GC-DZ). **(C)** Bar graph showing the mean shortest distance of CD57$^{lo}$ (black) and CD57$^{hi}$ (gray) TFH cells to DZ in individual follicles from one LN (left panel), and accumulated data (n=12 follicles from 5 LNs) showing the mean shortest distance of CD57$^{lo}$ (red) and CD57$^{hi}$ (green) TFH cells to Non-GC follicular area (right panel). **(D)** Plots showing CD4 T cell subsets from LNs (n=8) characterized by high expression of individual surface receptors or the expression of surface receptors per cell (judged by Mean Fluorescence Intensity, MFI) in cell subsets expressing a "receptor$^{hi}$" phenotype. **(E)** Plots showing the relative frequency of LN CD4 T cell subsets characterized by high expression of nuclear factors (n=8, LNs analyzed). **(F)** The intensity of each marker is shown (red-high expression and blue-low expression). 6000 PD-1$^{hi}$ memory CD4 T cells from each LN were analyzed. **(G)** tSNE analysis, based on the 9 clusters described in Fig. 4, for each individual LN analyzed. **(H)** The relative frequency of CD150$^{hi}$ (left) and CD151$^{hi}$ (right) CD4 T cell subsets from LNs (n=8) is shown. The Wilcoxon test was used for the analysis of data showed in C, D, E and H.

**Supplemental Figure 6. Analysis of synapse related receptors and sorting scheme of tonsillar CD4 T cell subsets. (A)** Flow cytometry plots showing the expression of CD151 and CD18 in total tonsillar CD4 T cells, naïve and memory CD4 T cell subsets. The red rectangles show the gating for the identification of receptor$^{hi}$ cells. **(B)** Graphs showing the relative frequency of tonsillar CD18$^{hi}$ and CD151$^{hi}$ CD4 T cell subsets (marked with different symbols) and the expression of active LFA-1 (MFI) in the same CD4 T cell subsets (n=10 tonsils). The expression of CD151 (MFI) in tonsillar CD4 T cell subsets marked with different symbols (n=10 tonsils) is also shown. The Wilcoxon test was used for the analysis of the data. **(C)** Schematic representation of supported lipid bilayer (SLB) tool. Artificial lipid bilayer is prepared on glass surface, to which anti-CD3 antibody, ICAM-1 and CD80 are incorporated via biotin or histidine tag. These proteins are highly mobile and support formation of highly organized immunological synapse upon interaction with T cells that sequester TCR and co-stimulatory molecules in the central supramolecular activation cluster (cSMAC) and integrin in the peripheral supramolecular activation cluster (pSMAC). **(D)** Bar graph showing the upregulation of CD69 in naïve CD4 T cells stimulated with SLB coated beads containing different concentrations of α-CD3, CD80 and ICAM-1. After 2 hr of stimulation, cells were analyzed for CD69 expression by flow cytometry. Arrow indicates the dose considered for most of experiments to study immunological synapse (lower panel) (n=2 repeats). **(E)** Flow cytometry plots showing the gating strategy for sorting relevant populations (left panel) and the co-expression of ICOS and PD-1 in TFH cells (right panel).

**Supplemental Figure 7. TIRF analysis of sorted tonsillar CD4 T cell subsets. (A)** Quantification of TCR microclusters using Icy software (Open BioImage). The "spread" image used for the calculation of the synaptic area (white line). **(B)** Graph showing the expression of actin in tonsillar CD4 T cell subsets (marked with different symbols) analyzed by flow cytometry (left panel) (n=5 tonsils). TIRF images showing the distribution of TCR and actin in the synapse of sorted CD4 T cell subsets (right panel). T cells were fixed after interaction with SLB for 20 min and stained for phalloidin-Alexa 647. Dot plots represent the area of actin depletion at the center of synapse (top) and the ratio of spread area of actin to the depletion zone (bottom) (n=3 tonsils). Accumulated data showing the average synaptic ICAM-1 (**C**) and TCR number of microclusters (**D**) in sorted populations (n=3 tonsils), marked with different color (ICOS$^{lo}$-black, ICOS$^{dim}$-blue and ICOS$^{hi}$-red). Each dot represents an individual cell. The Kruskal-Wallis and Conover-Inman methods were used for the statistical analysis. **(E)** The average synaptic TCR intensity in sorted populations, marked with different color (ICOS$^{lo}$-black, ICOS$^{dim}$-blue and CD57$^{lo}$ICOS$^{hi}$-green and CD57$^{hi}$ICOS$^{hi}$-red), from three tonsils is shown. Each dot represents an individual cell. The Welch ANOVA test was used for the analysis of presented data. Mean values (horizontal lines) and SD bars are shown.

**Supplemental Figure 8. Tissue imaging analysis of pTyr expression and TIRF analysis of sorted tonsillar TFH cells. (A)** Confocal image (40X, scale bar: 80 um) showing the expression of phospho-tyrosine (pTyr) staining and Ki67$^{hi}$ (top) in a tonsil tissue section (white circle depicts the borders of a follicle). The localization of CD20 (blue), CD3 (cyan), CD57 (red) and PD-1 (orange) positive cells are shown in a zoomed (scale bar: 40um) follicular area (middle panel). Merged images showing the expression of CD3-cyan/CD20-

blue/JoJo-gray, pTyr-green/JoJo-gray and pTyr-green/CD57-red/JoJo-gray in two zoomed areas (yellow circles) as well as at the interface of $CD57^{hi/lo}$ TFH cells interacting with B cells (bottom panel). **(B)** Confocal image showing specificity of pTyr staining in control vs λ-phosphatase treated cells. **(C)** Histocytometry plots showing the identification of non-follicular ($CD3^{hi}CD20^{lo}$) and follicular ($CD20^{hi/dim}Ki67^{hi}$) areas as well as the expression level of pTyr in $Ki67^{hi}$ vs $ki67^{low}$ B cells. **(D)** Flow cytometry 2D plots showing the expression of PTEN vs PD-1 in total live tonsillar cells. Data from two tonsils as well as the FMO control are shown. **(E)** Digital interference contrast (DIC) and epifluorescence image showing engulfment of labeled anti-CD3 antibody at the synapse of T cell interacting with SLB coated bead. **(F)** Bar graph showing the *ex vivo* proteasome activity in cell extracts derived from the indicated sorted CD4 T cell subsets (n=5 tonsils). **(G)** Dot plot showing the number of synaptic TCR microclusters in sorted tonsillar TFH cells (n=6 tonsils) treated either with DMSO (green) or MG132 (red) (right panel). The Welch ANOVA method was used for data analysis.
