## Supplementary figures and images for "Human follicular CD4 T cell function is defined by specific molecular, positional and TCR dynamic signatures"

### Supplementary figure 1

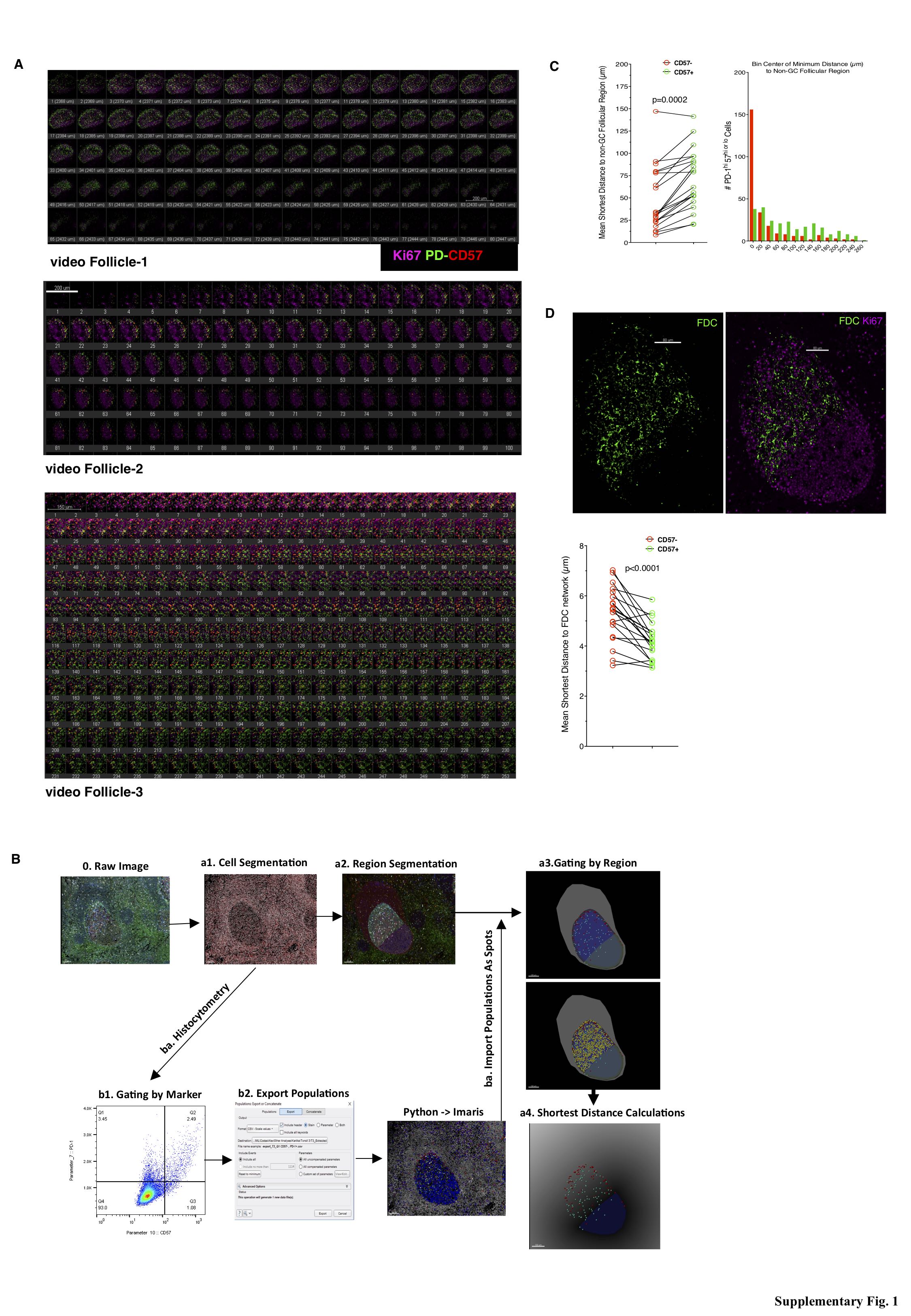

### Supplementary figure 2

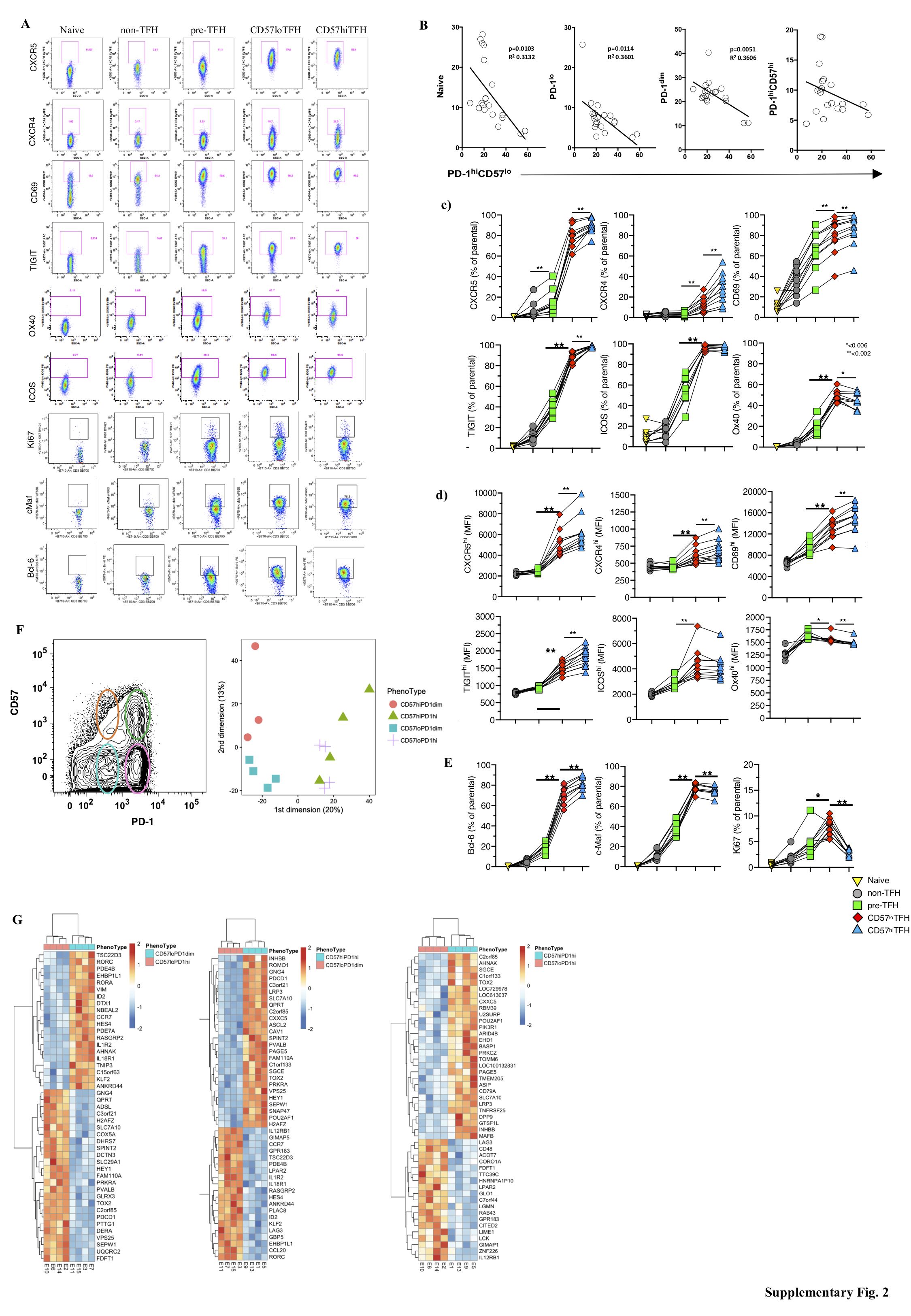

### Supplementary figure 3

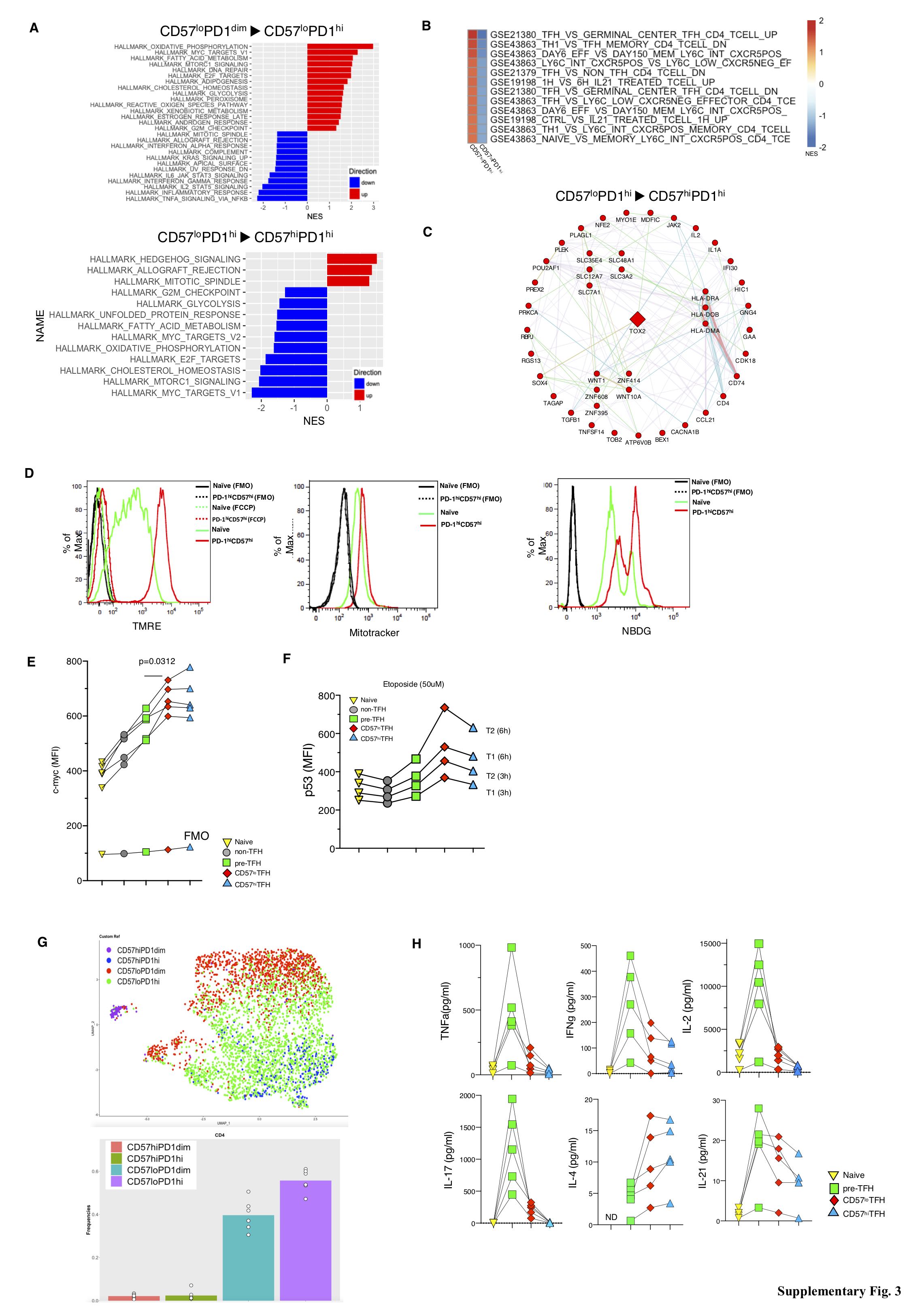

### Supplementary figure 4

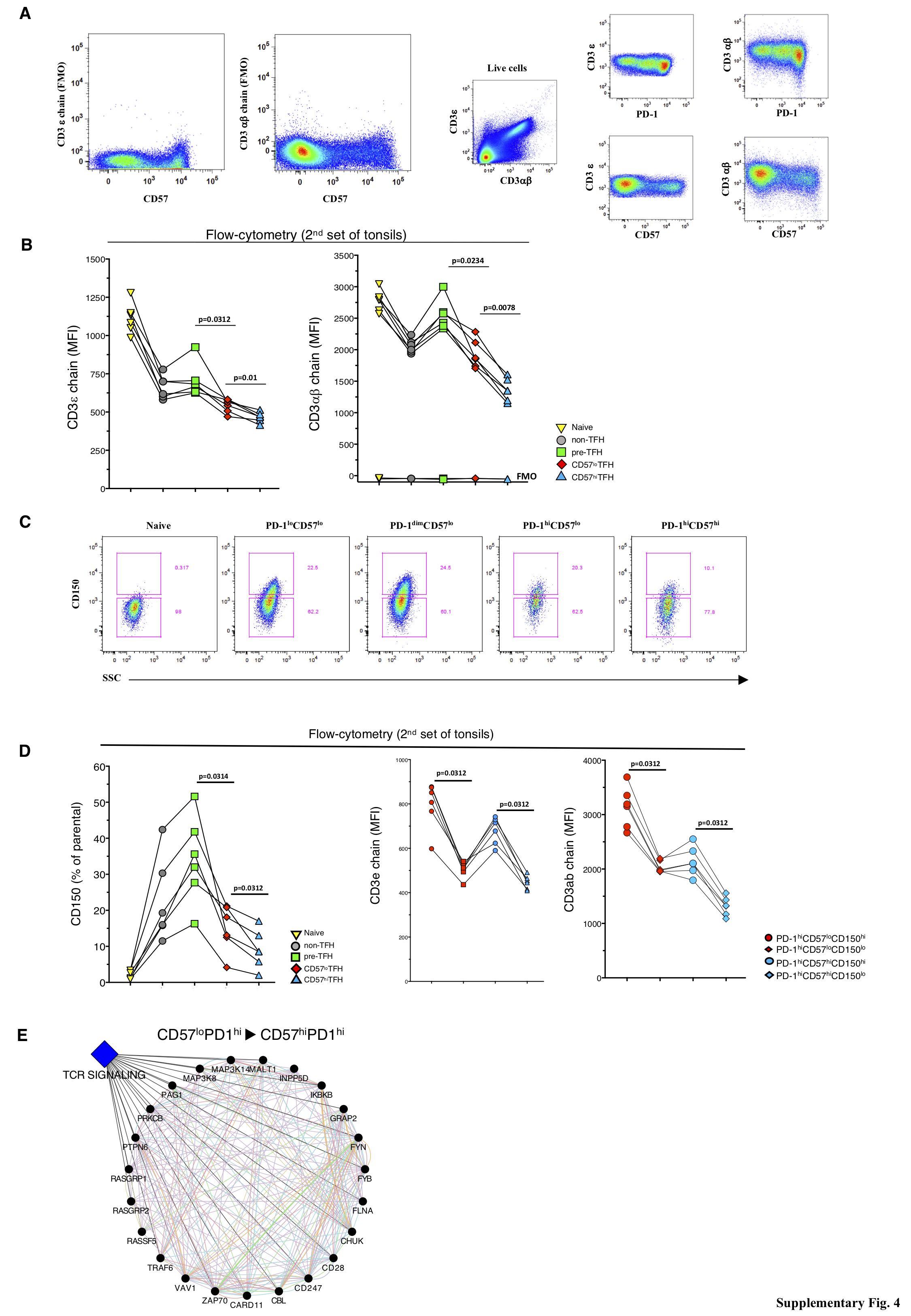

### Supplementary figure 5

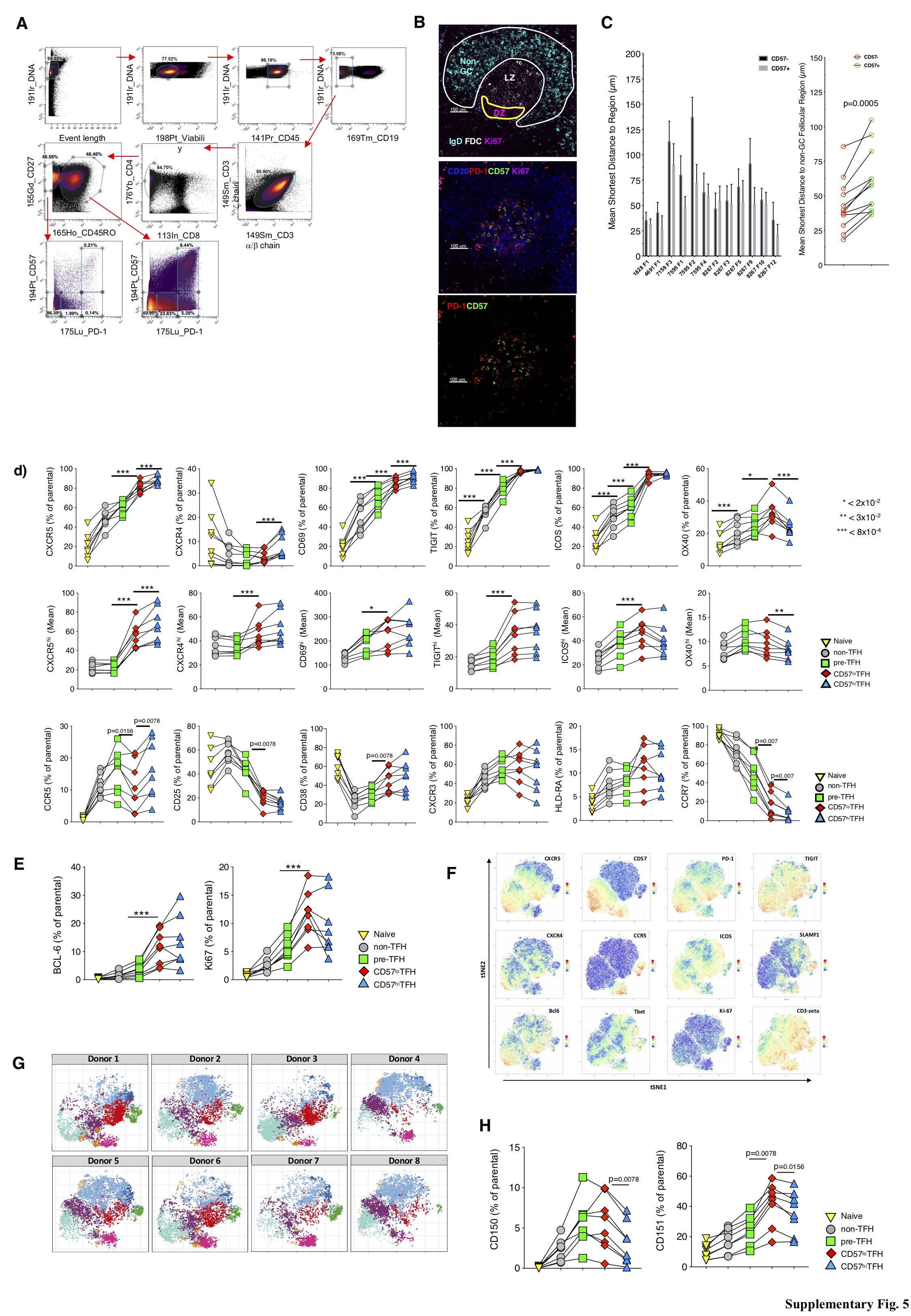

### Supplementary figure 6

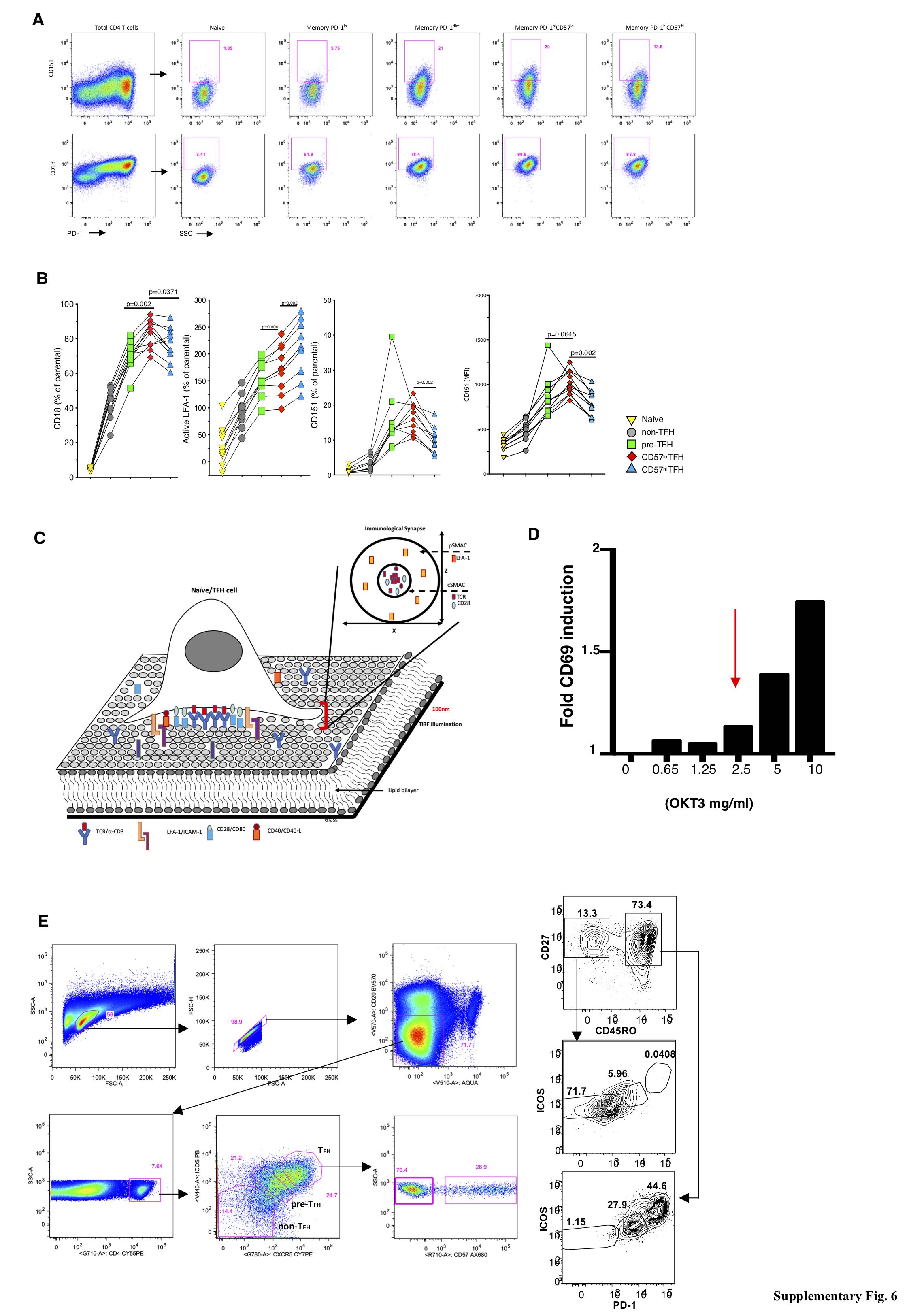

### Supplementary figure 7

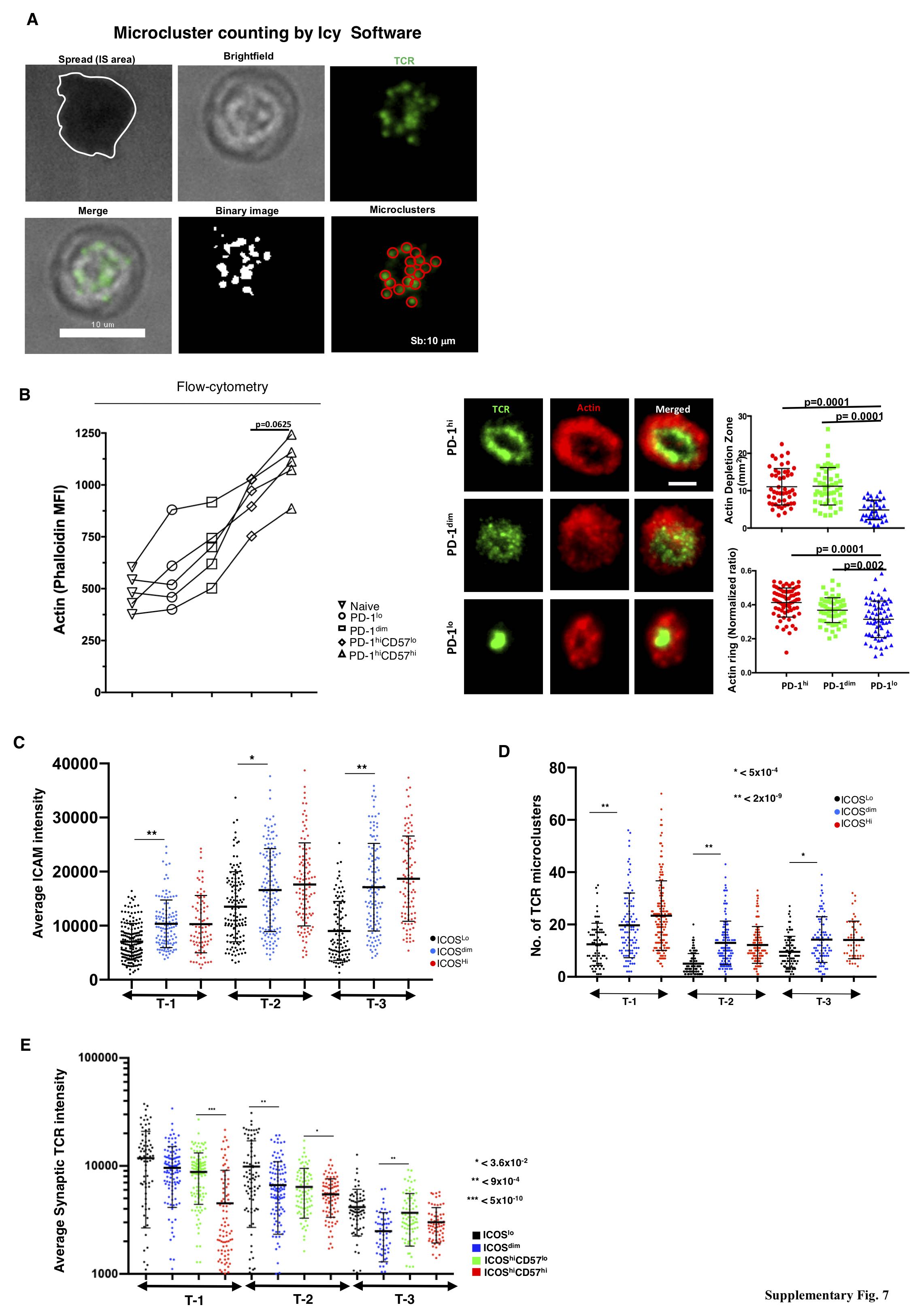

### Supplementary figure 8

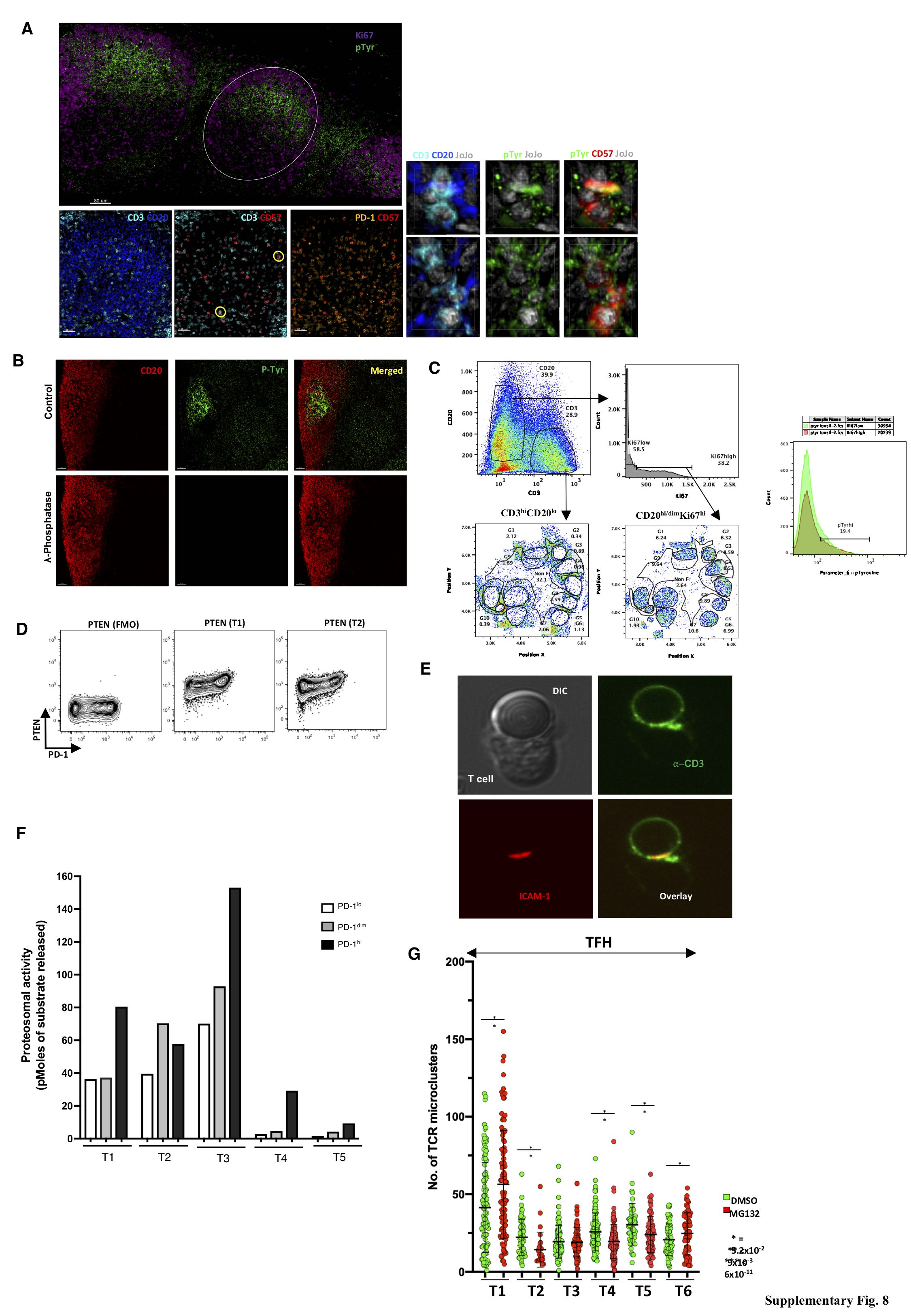
