## Supplementary movies legend for "Human follicular CD4 T cell function is defined by specific molecular, positional and TCR dynamic signatures"

**Supplementary Videos**

**Videos 1, 2 and 3:** 3D imaging of three tonsillar follicular areas is shown (CD20-cyan, Ki67-magenta, PD-1-green and CD57-red). The corresponding individual imaged planes are shown in the supplementary figure 3a.

**Videos 4,5 and 6:** live recording of *in vitro* immunological synapse formation. Sorted tonsillar CD4 T cells (PD-1^lo^, PD-1^dim^ and PD-1^hi^ TFH cells) were used (ICAM-red, TCR-green).
